# Sex-specific regulation of metabolic health and vertebrate lifespan by AMP biosynthesis

**DOI:** 10.1101/2022.01.10.475524

**Authors:** Gwendoline Astre, Tehila Atlan, Uri Goshtchevsky, Kobi Shapira, Adi Oron-Gottesman, Tomer Levy, Ariel Velan, Margarita Smirnov, Joris Deelen, Erez Y Levanon, Itamar Harel

**Affiliations:** Department of Genetics, the Silberman Institute, the Hebrew University of Jerusalem, Givat Ram, Jerusalem, 91904, Israel; Mina and Everard Goodman Faculty of Life Sciences, Bar-Ilan University, Ramat-Gan, 52900, Israel; Central Fish Health Laboratory, Department of Fisheries and Aquaculture, Ministry of Agriculture and Rural Development, Nir David, 10803, Israel; Max Planck Institute for Biology of Ageing, Cologne, Germany.; Molecular Epidemiology, Department of Biomedical Data Sciences, Leiden University Medical Center, Leiden, The Netherlands

**Author notes:** Equal contribution (Gwendoline Astre, Tehila Atlan).

## Abstract

Energy homeostasis is disrupted with age, which then fuels multiple age-related pathologies. The AMP-activated protein kinase (AMPK) is the primary sensor of cellular energy in eukaryotes. However, the genetic regulation of vertebrate aging by AMPK remains poorly understood. Here, we manipulate energy levels in the turquoise killifish by mutating *APRT*, a key enzyme in AMP biosynthesis. These manipulations produced a male-specific lifespan extension and restored metabolic plasticity. Exploring the observed sex differences using an integrated omics approach implicated the mitochondria as an important player. Mechanistically, APRT regulated mitochondrial functions and AMPK activity, mimicking energy starvation in heterozygous cells. A fasting-like state was also detected, particularly in heterozygous males, which leads to resistance to high-fat diet. Finally, life-long intermittent fasting eliminated the male-specific longevity benefits mediated by the *APRT* mutation. These observations identify the AMP/AMPK axis as a sex-specific regulator of vertebrate longevity and metabolic health.

## Introduction

Aging is associated with metabolic changes, including deregulated nutrient-sensing and mitochondrial dysfunction (Amorim et al., 2022; Lopez-Otin et al., 2013). These changes are mechanistically linked to the onset of many age-related diseases, including diabetes, neurodegeneration, and cancer (Finkel, 2015; Niccoli and Partridge, 2012). Accordingly, extensive efforts have been invested in exploring pro-longevity interventions (either dietary, genetic, or pharmacological), by targeting primary metabolic pathways (Smith et al., 2020). However, many aspects of energy metabolism are differently regulated between males and females (Link and Reue, 2017; Mauvais-Jarvis et al., 2017; Wu and O’Sullivan, 2011), contributing to a sex-biased predisposition to age-related diseases (Hägg and Jylhävä, 2021; Tramunt et al., 2020). Thus, understanding the enigmatic molecular mechanisms that mediate sex differences in longevity is crucial for designing tailored interventions (Austad and Fischer, 2016).

In the context of vertebrate aging, a series of specialized nutrient sensing pathways have been genetically investigated in mice. These include the insulin and insulin-like growth factor-1 (IGF-1 pathway, which senses glucose levels), and the downstream mechanistic target of rapamycin (mTOR pathway, which senses amino acid levels) (Folgueras et al., 2018; Green et al., 2021; Johnson et al., 2013). Yet, surprisingly, the genetic regulation of vertebrate aging by AMP-activated protein kinase (AMPK), an evolutionary conserved sensor of cellular energy levels, remains poorly understood (Burkewitz et al., 2014; Green et al., 2021; Smith et al., 2020).

AMPK is activated in response to low energy levels, which are reflected by a higher AMP:ATP ratio in the cell. Subsequently, AMPK activation can restore the energy balance by inhibiting ATP-consuming processes, and activating ATP-generating processes (Garcia and Shaw, 2017; Hardie et al., 2012). Thus, AMPK is involved in many evolutionarily conserved aging-related interventions, such as dietary restriction, fasting, physical exercise, and metformin administration (Green et al., 2021; Smith et al., 2020). However, genetic mouse models with constitutively active AMPK have so far demonstrated conflicting, and even detrimental metabolic phenotypes (Carling, 2017; Garcia and Shaw, 2017; Woods et al., 2017; Yang et al., 2016; Yavari et al., 2016).

In contrast, genetic manipulation of the AMP/AMPK axis has been shown to extend the lifespan of invertebrate models (Apfeld et al., 2004; Curtis et al., 2006; Greer et al., 2007; Mair et al., 2011; Matecic et al., 2010; Stenesen et al., 2013; Ulgherait et al., 2014). However, as seen in *Drosophila* (Stenesen et al., 2013), the proposed mechanism in some cases was thought to be invertebrate-specific (Camici et al., 2018). Thus, the conservation of similar genetic networks in vertebrate aging remains unclear (Smith et al., 2020). Developing new genetic strategies will be required to better understand how the AMP/AMPK axis regulates vertebrate metabolism and longevity, either by tissue-specific manipulation of AMPK (see accompanying paper by Ripa et al., and (Garcia et al., 2019)), or by systemic manipulation of energy homeostasis, as explored in this study. So far, genetic dissection of vertebrate aging has been hindered by the relatively long lifespan of the classical mouse and zebrafish models (Harel and Brunet, 2015). Thus, developing alternative vertebrate platforms could significantly facilitate experimental approaches.

The African turquoise killifish *Nothobranchius furzeri* has recently emerged as a powerful vertebrate model (Astre et al., 2022; Baumgart et al., 2016; Cellerino et al., 2015; Harel and Brunet, 2015; Kim et al., 2016). These fish provide an attractive experimental platform as they possess a naturally compressed lifespan that is 6 or 10 times shorter than mice or zebrafish, respectively (Tacutu et al., 2018). Killifish exhibit typical vertebrate aging phenotypes, including a decline in cognitive functions and an increase in age-related pathologies (Astre et al., 2022; Cellerino et al., 2015; Harel and Brunet, 2015; Kim et al., 2016), and they respond to conserved lifespan interventions (including drug treatments and dietary restriction (Terzibasi et al., 2009; Valenzano et al., 2006)). The recently developed integrative genomic and CRISPR-based genome-editing platform has transformed the killifish into a bona fide genetic model for rapid exploration of vertebrate aging and diseases (Astre et al., 2022; Harel et al., 2015, 2016; Reichwald et al., 2015; Valenzano et al., 2015). However, genetic exploration of pro-longevity pathways has not been attempted so far.

Here we leverage the turquoise killifish as a powerful model to explore the genetic regulation of vertebrate aging by the energy-sensing AMP/AMPK axis. In order to manipulate ATP levels, we generate a heterozygous mutation in the *APRT* gene, a key enzyme in AMP biosynthesis. AMP, the precursor of ADP and ATP, is formed by either the *de-novo* or salvage AMP biosynthetic pathways (Camici et al., 2018). The AMP salvage pathway is more energy efficient, and is predicted to be the primary source (∼90%) of daily purine nucleotide biosynthesis across different organs (Johnson et al., 2019; L Ipata et al., 2011; Murray, 1971; Young et al., 2021). We then demonstrate that the *APRT* mutation extends the lifespan in a male-specific manner, and restores metabolic plasticity in old age (including glucose homeostasis). Interestingly, our findings are so far specific to APRT, as mutating another enzyme in the ATP synthesis pathway has no effect on the lifespan.

We further identified a distinct gene expression signature that links mitochondrial numbers and functions with the sex differences in longevity, and have developed a cell-culture platform for killifish that allows us to explore the mechanisms involved. We use this system to show a reduction in mitochondrial functions and ATP levels, and an increase in AMPK activity in cells derived from *APRT* mutant males, which display a phenotype that resembles a persistent state of starvation. These results are supported by a fasting-like state and a resistance to a high-fat diet, seen exclusively in male mutants, and a functional role of sex-hormones in altering mitochondrial biogenesis. Finally, intermittent fasting eliminates the sex-specific longevity benefits mediated by the *APRT* mutation. Collectively, these data identify the AMP salvage pathway as a novel genetic regulator of vertebrate aging, and APRT as a promising pharmacological target for pro-longevity interventions.

## Results

### Generation of a genetic model for APRT, a member of the AMP biosynthesis pathway

To genetically alter organismal energy homeostasis, we manipulated AMP biosynthesis (Figures 1, S1). For this purpose, we selected adenine phosphoribosyltransferase (APRT), which catalyzes the formation of AMP from adenine and PRPP (5-phosphoribosyl-1-pyrophosphate) as our primary target (Figure 1a). CRISPR-based genome editing (Astre et al., 2022), was used to generate an 8 base-pair deletion (Δ8) allele in the killifish *APRT* gene, which was predicted to give rise to a premature stop codon and a loss of function (Figure 1b). We then outcrossed this line for 8 generations to reduce the effect of potential off-target mutations.

**Figure 1:**
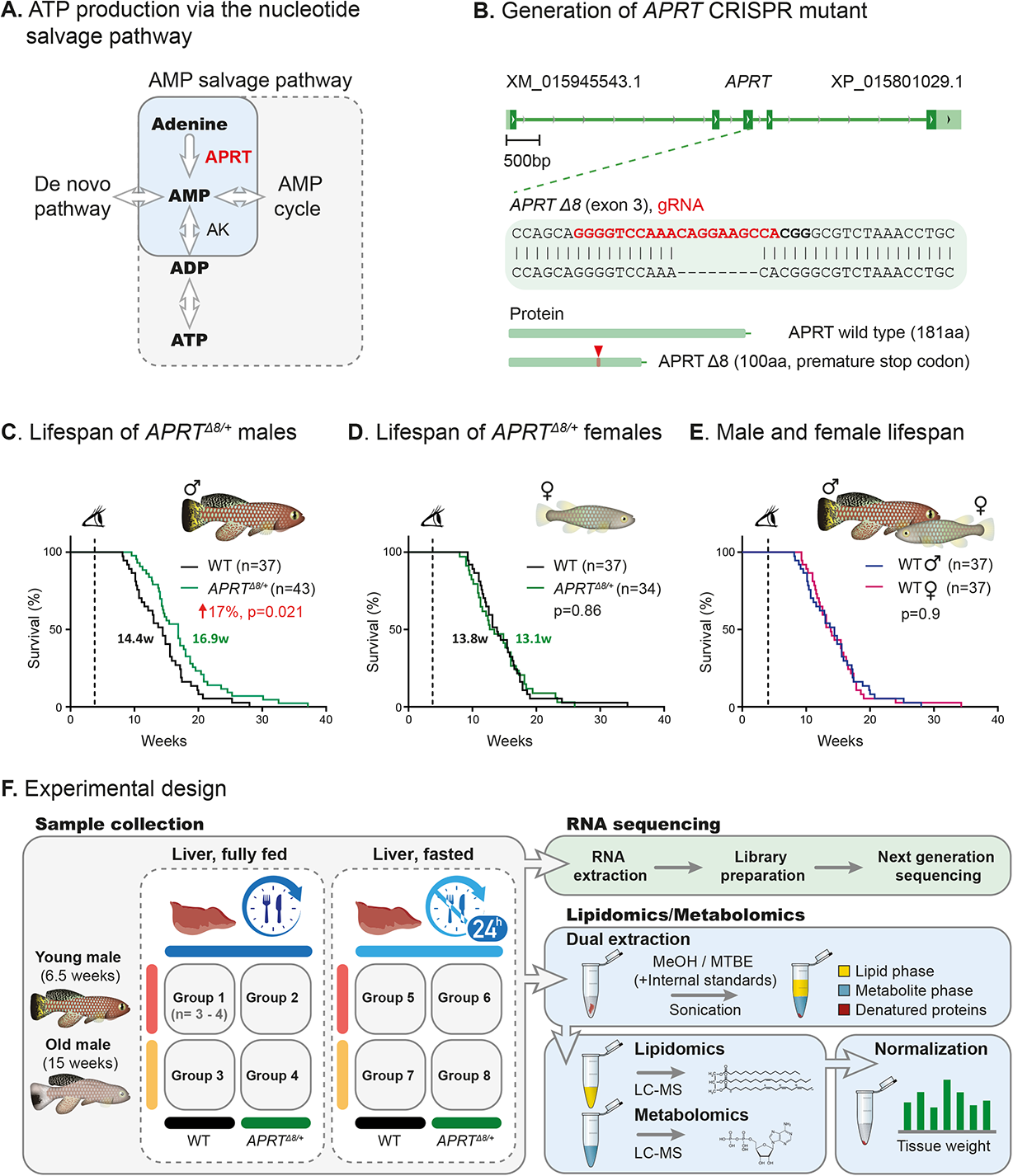
Male-specific lifespan extension by targeting *APRT*, a key enzyme in the AMP biosynthesis. (**A**) As a part of the nucleotide salvage pathway, the enzyme adenine phosphoribosyltransferase (APRT) catalyzes adenine into AMP, a key precursor for ATP production. AMP can be also produced by other pathways, including the AMP *de*-*novo* pathway and the AMP cycle. adenylate kinase (AK) enzymes catalyze the reversible conversion of adenine nucleotides (2ADP = ATP + AMP) to generate ATP. (**B**) Generation of *APRT* CRISPR mutant, including guide RNA (gRNA) target for exon 3 (red sequence), protospacer adjacent motif (or PAM, in bold), and successful germline transmission of an 8 bp deletion (Δ8). The *APRT* Δ8 allele is predicted to generate a protein with a premature stop codon. (**C-E**) Lifespan of WT and *APRT^Δ8/+^* fish assessed separately for males (C) and females (D). (E) Overlay of the male and female lifespans from WT fish, using data from (C) and (D). p-values for differential survival in log-rank tests, median survival, and fish numbers are indicated. See Table S1 for additional information. (**F**) Analysis pipeline for characterizing the livers of male fish using RNA sequencing, metabolomics, and lipidomics. Experimental groups differ by age (young or old), genotype (WT or *APRT^Δ8/+^*), and feeding condition (fully-fed or fasted for 24 h).

Mating heterozygous *APRT^Δ8/+^* pairs follows the expected Mendelian ratios with respect to the genotypes of the fertilized eggs (p = 0.89, χ2 test, Figure S1a), and there was no significant difference in either fecundity of the pairs (Figure S1b), or the size of produced eggs (Figure S1c). However, after hatching, we failed to detect any adult *APRT^Δ8/Δ8^* fish, suggesting that the homozygous mutation causes embryonic lethality (p = 0.0006, χ^2^ test, Figure S1a). Notably, an *APRT* homozygous mutation is associated with premature death after weaning in mice (Engle et al., 1996)), and embryonic lethal in *Drosophila* (Stenesen et al., 2013). In view of the central role of AMP biosynthesis in energy homeostasis, we were curious to examine the effect of the *APRT* heterozygous mutation on vertebrate lifespan.

### Male-specific lifespan extension in *APRT^Δ8/+^* heterozygous fish

Our results indicate that *APRT^Δ8/+^* heterozygous male fish live significantly longer (17% increase, log-rank test, p = 0.021, Figure 1c). In contrast, the female lifespan is not significantly affected (log-rank test, p = 0.86) (Figure 1d), and remains comparable to that of wild-type males (Figure 1e). In order to examine the specificity of our phenotype, we also mutated adenylate kinase 2 (AK2), a second enzyme in the ATP synthesis pathway (Stenesen et al., 2013). AK2, like other AK isozymes, catalyzes the reversible conversion of the AMP and ADP adenine nucleotides (2ADP = ATP + AMP) (Camici et al., 2018) (Figures 1a, S1d). A ‘knock-in” approach (Astre et al., 2022), was used to generate an *AK2^Δ8/+^* allele (Figure S1e).

Similar to the *APRT* mutation, the *AK2^Δ8/Δ8^* homozygous mutation is embryonic lethal (Figure S1f), But, in contrast to the findings in invertebrate models (Stenesen et al., 2013), no lifespan extension was observed in either male (log-rank test, p = 0.87) or female (log-rank test, p = 0.7) heterozygous fish (Figure S1g,h). Taken together, our findings highlight the role of the AMP salvage pathway, and specifically of the APRT enzyme, in modulating vertebrate lifespan. Furthermore, our sex-specific data provide an exciting opportunity to explore the molecular mechanisms of sex differences in aging (Austad, 2006; Austad and Fischer, 2016; Horstman et al., 2012). To examine how the longevity benefits by the *APRT^Δ8/+^* mutation are produced, we performed an integrated multi-omics analysis, initially focusing on male fish (Figure 1f).

### *APRT* heterozygous fish display an age-dependent signature of canonical AMPK targets

Energy homeostasis and metabolic functions depend on organismal age and nutrient availability (Lopez-Otin et al., 2013). We therefore performed a transcriptomic analysis comparing the liver, a primary metabolic organ, from young (6.5 weeks) or old (15 weeks) male wild-type (WT) or *APRT^Δ8/+^* heterozygous fish. To evaluate the influence of food availability and metabolic plasticity, we compared the livers from fully fed or fasted animals (fasting for 24 h) of each age/genotype, in a total of 8 experimental groups (Figure 1f). This scheme was designed to examine whether nutrient sensing is indeed deregulated during killifish aging, and if so, to determine the effect of the *APRT^Δ8/+^* mutation.

A principal component analysis (PCA) for transcript levels revealed separation of samples depending on the experimental conditions (Figure 2a for PC1/3, and Figure S2a for PC1/2). For example, old *APRT^Δ8/+^* heterozygous fish separated with young fish, suggesting a possible rejuvenated phenotype (Figure 2a, top). Age-dependent (Figure 2a, bottom) and genotype-dependent (Figure S2a, left) patterns were also observed. As expected, the levels of *APRT* transcripts were significantly reduced in heterozygous fish, probably as a result of nonsense-mediated mRNA decay (NMD) (Figure S2b). These findings further suggest that the *APRT^Δ8^* is a loss-of-function allele (there are currently no available antibodies directed against the killifish APRT protein).

**Figure 2:**
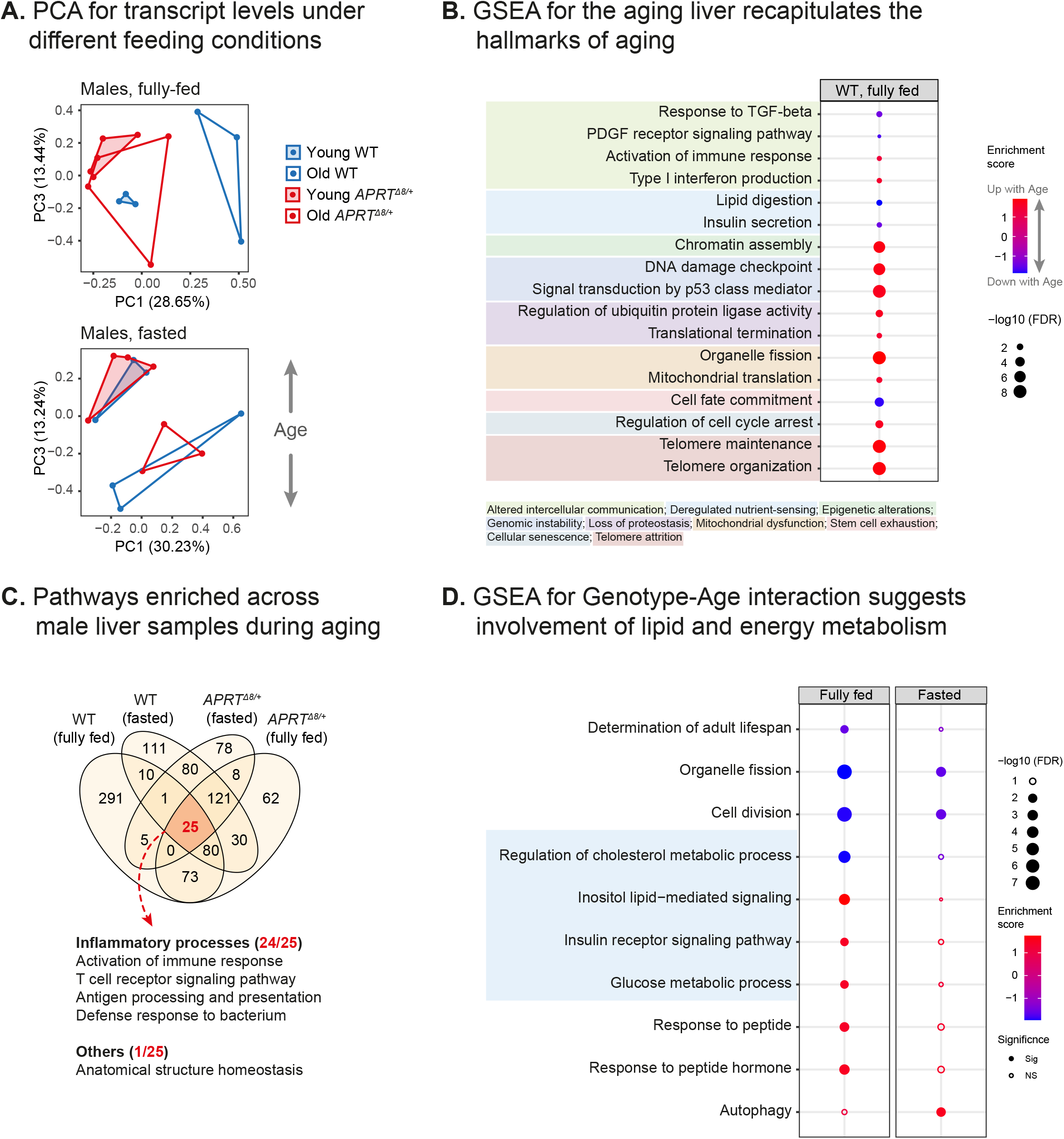
Transcriptomic analysis in males suggests a modified energy and lipid metabolism in the livers of old *APRT^Δ8/+^* fish. (**A**) Principal component analysis (PCA) for transcript levels from the livers of WT (blue) or *APRT^Δ8/+^* (red) fish, either young (shaded) or old (nonshaded). Experiments were performed under fully-fed (top) or fasting conditions (bottom). Each symbol represents an individual fish. PC1/PC3 are shown. PC1/PC2 are reported in Figure S2. (**B**) Functional enrichments (GO) using gene set enrichment analysis (GSEA) for differential RNA expression with aging (WT, fully-fed). Enrichment significance was at FDR < 5%. (**C**) Venn diagram showing the overlap of significantly enriched pathways with aging across all male samples. GO enrichments using GSEA were called at FDR < 5%. (**D**) Genotype-age interaction analysis to identify enriched pathways (using GSEA) that co-depend on both genotype and age (e.g. how the *APRT^Δ8/+^*mutation modifies the aging process). Analysis was performed separately for fully-fed or fasting conditions. GO Enrichments using GSEA were called at FDR < 5%.

In order to obtain an overview of hepatic aging in killifish, we first screened for genes differentially expressed between young and old control fish (WT, fully fed), and conducted gene set enrichment analysis (GSEA) using gene ontology (GO). The results revealed 1242 downregulated and 1509 upregulated genes, with an FDR of 0.05. Among the significantly enriched pathways, we could identify classical ‘hallmarks of aging’ signatures, including genes associated with genomic instability, mitochondrial dysfunction, deregulated nutrient sensing, and cell cycle (Figure 2b, Table S2). To identify a robust signature of aging, we then expanded this analysis to include all age-related changes, independent of genotype or feeding condition (Figures 2c, S2c). This comparison identified increases in inflammatory responses, or ‘inflammaging’ (Benayoun et al., 2019; Franceschi et al., 2018), as a robust age-related transcriptional signature. Similarly, a comprehensive histological examination of selected organs highlighted age-dependent degeneration of the kidneys, a primary lymphoid organ in fish (Figure S2d). See Table S1 for summary of pathological findings (n≥3 for each group).

Are old *APRT^Δ8/+^* fish truly rejuvenated, as predicted by the PCA? To identify differentially expressed genes that co-depend on both genotype and age, we performed interaction analysis, followed by GSEA using GO. Under both feeding conditions, we identified canonical AMPK downstream targets that are co-dependent on both age and genotype (Figure 2d, Table S2). These downstream targets are involved in autophagy, energy metabolism, (i.e. insulin and glucose homeostasis) and lipid metabolism, as well as determination of adult lifespan, and cell-cycle-related pathways.

### Age-related decline in nutrient sensing is prevented in old *APRT^Δ8/+^* fish

To further explore energy metabolism, we performed metabolomic analysis on killifish livers (Figures 3, S3a), using the same experimental design performed for the transcriptomics (Figure 1f). For this purpose, we focused on polar metabolites, such as nucleic acids, sugars, and small organic acids that are typically involved in primary metabolism. Similar to the transcriptomic analysis (Figure 2), PCA demonstrated that under fully fed conditions, samples primarily segregated according to age (Figure 3a, top), while genotype dependency was revealed under fasting conditions (Figure 3a, bottom). These data confirm the importance of feeding regimens in exposing age- and genotype-specific metabolic signatures.

**Figure 3:**
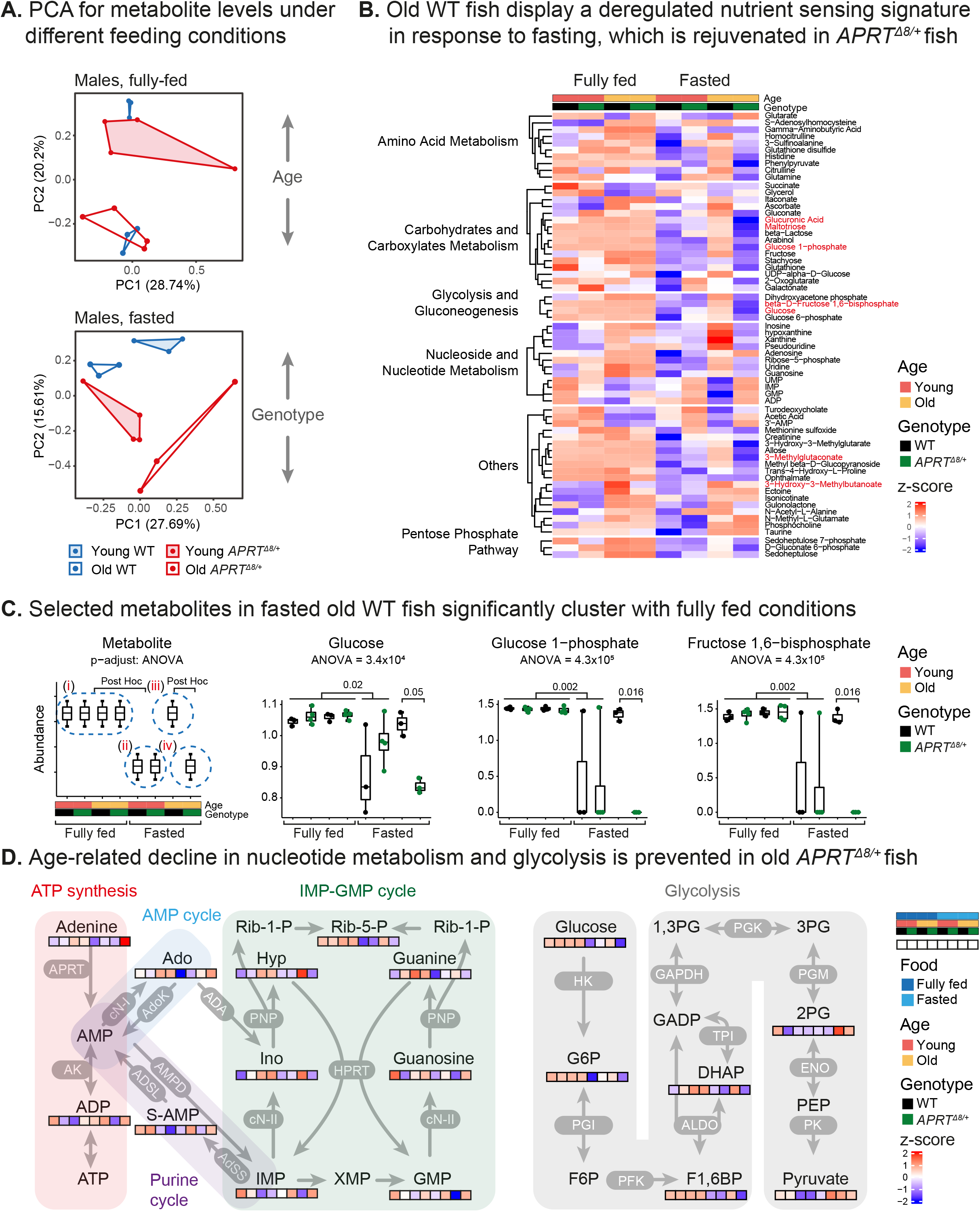
Metabolomic analysis demonstrates restoration of energy homeostasis in livers of old *APRT^Δ8/+^* fish. (**A**) Principal component analysis (PCA) for metabolite levels from the livers of WT (blue) or *APRT^Δ8/+^* (red) fish, young (shaded) or old (nonshaded). Experiments were performed under fully-fed (top) or fasting conditions (bottom). Each symbol represents an individual fish. (**B**) Heatmap showing significantly altered liver metabolites clustered into key metabolic pathways (full metabolite list is reported in Figure S3 and Table S3). Each square represents the normalized relative abundance (indicated by color), from an average of 3-4 fish. Significant metabolites were evaluated by one-way ANOVA between 4 clusters as depicted in (C): fully fed (**i**); young fasting (**ii**); old WT fasting (**iii**); and old *APRT^Δ8/+^* fasting (**iv**). Using these clusters, we can estimate the normal response to fasting (comparing clusters **i** and **ii**), and detect the age-dependent deregulated nutrient sensing that is rejuvenated in old *APRT^Δ8/+^* fish (comparing clusters **iii** and **iv**). Significance was called at FDR ≤ 5% for both ANOVA and Tukey post-hoc. The full statistical values are reported in Table S3. (**C**) Boxplot showing significant metabolites selected from (B), of carbohydrate and glucose metabolism. n = 3-4 fish per experimental condition, each symbol represents one fish. Bars represent minimum and maximum. (**D**) Selected metabolic pathways depicting key enzymes and abundance levels of main compounds. Each square represents the color-coded normalized relative abundance as seen in (B). Pathways were curated from the KEGG (Kanehisa, 2017) and BioCyc (Caspi et al., 2020). 2PG: 2-Phospho-D-Glycerate; Ado: Adenosine; DHAP: Dihydroxyacetone phosphate; F1,6BP: D-Fructose 1,6-bisphosphate; G6P: Glucose 6-phosphate; Hyp: hypoxanthine; Ino: Inosine; Rib-5-P: Ribose-5-phosphate; S-AMP: Adenylo-Succinate; Rib-1-P: Ribose-1-phosphate; F6P: Fructose 6-phosphate; 1,3PG: 1,3-Bisphosphoglycerate; GADP: Glyceraldehyde 3-phosphate; 3PG: 3-Phosphoglyceric acid; PEP: Phosphoenolpyruvate

Out of a total of 160 unique metabolites, 140 passed our quality control tests (Table S3). We then applied hierarchical clustering to all 140 detected metabolites, and focused on a subgroup whose signature displays the following characteristics (Figure S3a): **a)** dependent on nutrient availability (i.e. food dependent); **b)** exhibits significant differences between experimental conditions; **c)** displays a restoration of youthful levels in old heterozygous fish under fasting conditions (see Methods for full statistical analysis). These selected metabolites are members of primary energy and biosynthetic pathways, including amino acid and carbohydrates metabolism, glycolysis, nucleoside and nucleotide metabolism, and the pentose phosphate pathway (PPP) (Figure 3b,c).

For example, following fasting, the levels of PPP-related metabolites were reduced in the livers of young fish and old *APRT* heterozygous fish, while remaining relatively high in old WT fish. Limiting the PPP is an integral part of the mammalian response to fasting (Adams et al., 1983; Jenniskens et al., 2002; Lee et al., 2019; Meister, 1988), and inhibition of the PPP was recently shown to promote longevity in *C. elegans* by mimicking dietary restriction (Bennett et al., 2017). Overlaying the levels of individual metabolites onto the glycolysis and ATP salvage pathways, demonstrated a similar behavior (e.g. Fructose 1,6−bisphosphate, Figure 3c,d), with some metabolites displaying alternative patterns (e.g. GMP and IMP). Taken together, our metabolomic analysis suggests that old fish suffer from deregulated nutrient sensing, which is partially rejuvenated in old *APRT^Δ8/+^*heterozygous fish.

The genotype-age interactions also predicted altered lipid metabolism (Figure 2d), which has been recently implicated in aging and longevity in both mammals (Johnson and Stolzing, 2019; Palavicini and Han, 2021) and fish (Ahuja et al., 2019; Milinkovitch et al., 2018). This prompted the question as to which lipid species are linked to the metabolic rejuvenation observed in old *APRT^Δ8/+^* fish. Similar to the metabolomic signatures, the PCA demonstrated age-dependent differences under fully-fed conditions (Figure S3b, top), with a genotype contribution when the animals were fasted (Figure S3b, bottom). Applying hierarchical clustering, revealed a complex signature that is affected by age, genotype, and feeding conditions (Figure S3c, left). We then scanned our data for lipid species that exhibit restoration of youthful patterns in old *APRT^Δ8/+^* heterozygous fish under fasting conditions. The results highlighted a distinct group of glycerophospholipids, including lysophosphatidylcholine (LPC), lysophosphatidylethanolamine (LPE), and phosphatidylinositols (PI) (Figure S3c,d).

Interestingly, while the level of LPEs increased with age during fasting in WT animals, the heterozygous old fish retained the lower levels characteristic of youth. In this context, low levels of plasma LPE (i.e. LPE(P-16:0)) were recently associated with exceptional human longevity (Pradas et al., 2019), and LPC levels have been implicated in human aging and mitochondrial functions (Semba et al., 2019). It may also be pertinent that although PIs constitute only a small portion of cellular phospholipids, they have recently emerged as key modulators of energy metabolism, including insulin sensitivity and obesity (Balla, 2013; Bridges and Saltiel, 2015; Lundquist et al., 2018). Taken together, our findings highlight a role for the AMP salvage pathway, and specifically for the APRT enzyme, in modulating vertebrate lifespan and restoring metabolic plasticity in old males.

### Genotype-age interaction analysis identifies a male-specific mitochondrial signature

What mediates the observed sex differences? In order to address this question we performed transcriptomic analysis on *APRT^Δ8/+^* heterozygous females under metabolic baseline (experimental groups #9-12, Figure 4a, S4a). In agreement with the findings in males, *APRT* transcripts were significantly reduced in heterozygous fish (Figures S2b, S4b). GSEA for age-related differential gene expression confirmed that ‘inflammaging’ is a conserved aging hallmark in both sexes (Figure 4b, Table S2). However, there were distinct sex-linked biases observed, specifically with respect to carbohydrate, steroid, and lipid metabolism (Figure 4b,c).

**Figure 4:**
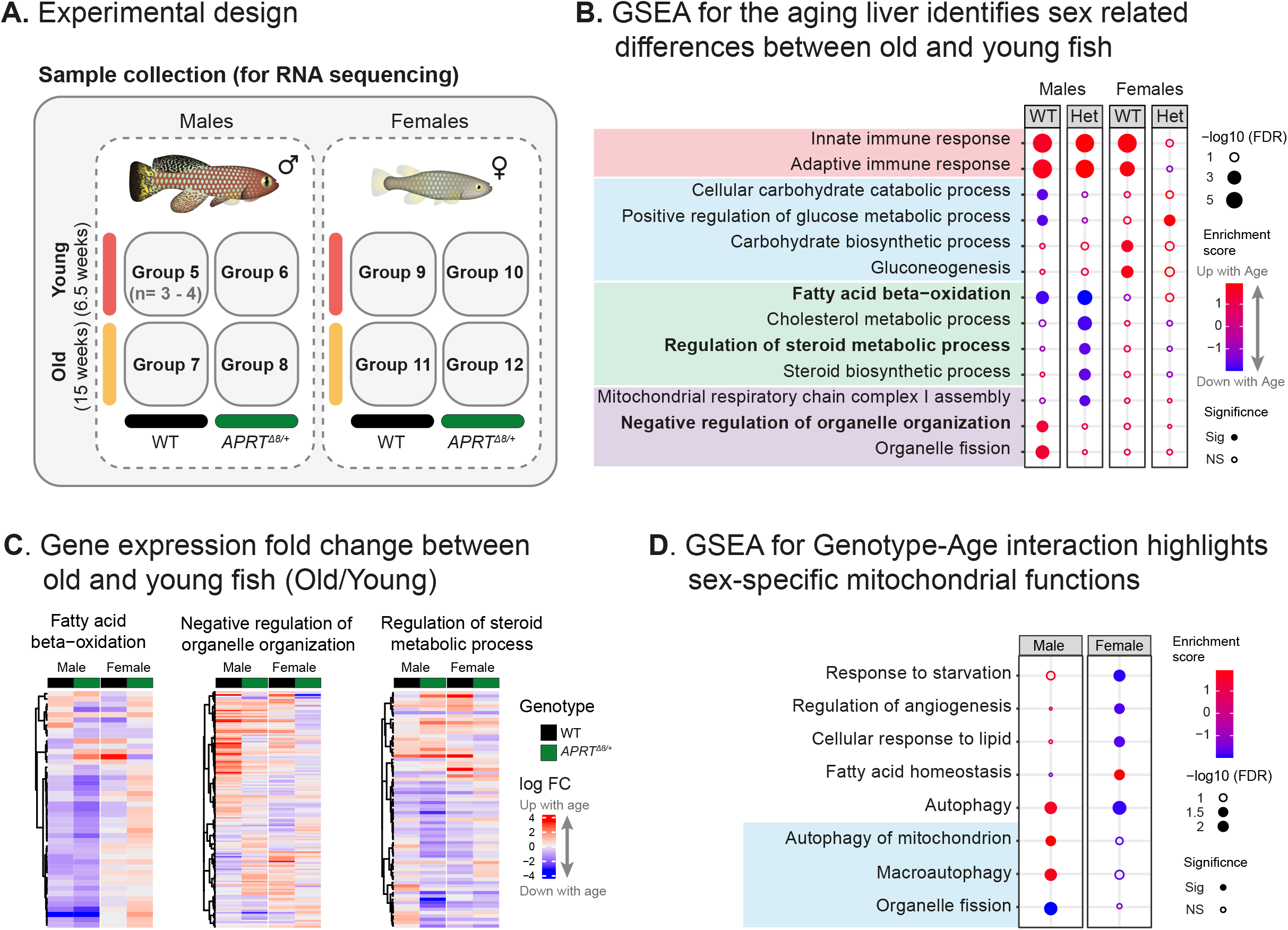
The male-specific lifespan extension in *APRT^Δ8/+^* fish is associated with a distinct mitochondrial transcriptional signature. (**A**) Experimental design for characterizing the livers of male and female fish using RNA sequencing. Experimental groups differ by age (young or old) and genotype (WT or *APRT^Δ8/+^*). Note that there is a single feeding condition in this design (i.e. 24 h fasting), and that male fish (experimental groups #5-8) are also presented in Figure 1f. (**B**) Functional enrichments (GO) using gene set enrichment analysis (GSEA) for differential RNA expression between old and young fish. A comparison between the different experimental groups demonstrates conserved (e.g., inflammatory processes, in red) and sex-specific dynamics. Sex-specific pathways include energy metabolism, specifically carbohydrate and lipid metabolism (in blue or green, respectively) and mitochondrial functions (in purple). Pathways in bold are also depicted in (C). GO enrichments using GSEA were called at FDR < 5%. (**C**) Heatmap for age-related gene expression changes in male and female livers (representative pathways are selected from (B)). Genes are hierarchically clustered, and expression values are presented as log2 (fold-change) between old and young using a 1.5-fold change cutoff. (**D**) Genotype-age interaction analysis was used to identify sex-specific pathways that co-depend on both genotype and age. Functional enrichments (GO) were performed using gene set enrichment analysis (GSEA). GO enrichments using GSEA were called at FDR < 5%.

Since, we anticipated that pathways modified particularly in male *APRT^Δ8/+^* fish during aging, are more likely to make a functional contribution to the longevity effect, we performed a similar genotype-age interaction analysis on the female transcriptomic data (Figure 2d). Comparing our findings between sexes indeed highlighted several sex-dependent pathways, including an altered response to starvation, as well as several male-specific mitochondrial-related pathways (Figure 4d, Table S2). However, as the interaction analysis considers both genotype and age, how the largely opposite responses between sexes are reflected in gene expression is non-intuitive. We therefore calculated the fold change between the gene expression of heterozygous and WT fish for each experimental condition, which provided a simple representation of the sex-biased transcriptional response (Figure S4c).

The result of this analysis predicts that mitochondrial dynamics, such as copy numbers and function, could be co-dependent on age, sex, and genotype. Among the core regulators of mitochondrial dynamics, genes responsible for mitochondrial biogenesis displayed a strong sex-dependency when compared to fission or fusion (Figure S4d). This behavior is particularly observed in *PGC-1α* (peroxisome proliferator-activated receptor gamma coactivator 1-α), the master regulator of mitochondrial biogenesis, and is consistent with AMPK activation (i.e. *PGC-1α* is an AMPK target (Garcia and Shaw, 2017)). Accordingly, a significant increase in mitochondrial DNA (mtDNA) copy number was detected, specifically in the livers of old male *APRT^Δ8/+^* fish (Figures 5a). Similar patterns were observed in other organs, such as the tail, suggesting a systemic response (Figure 5b).

**Figure 5:**
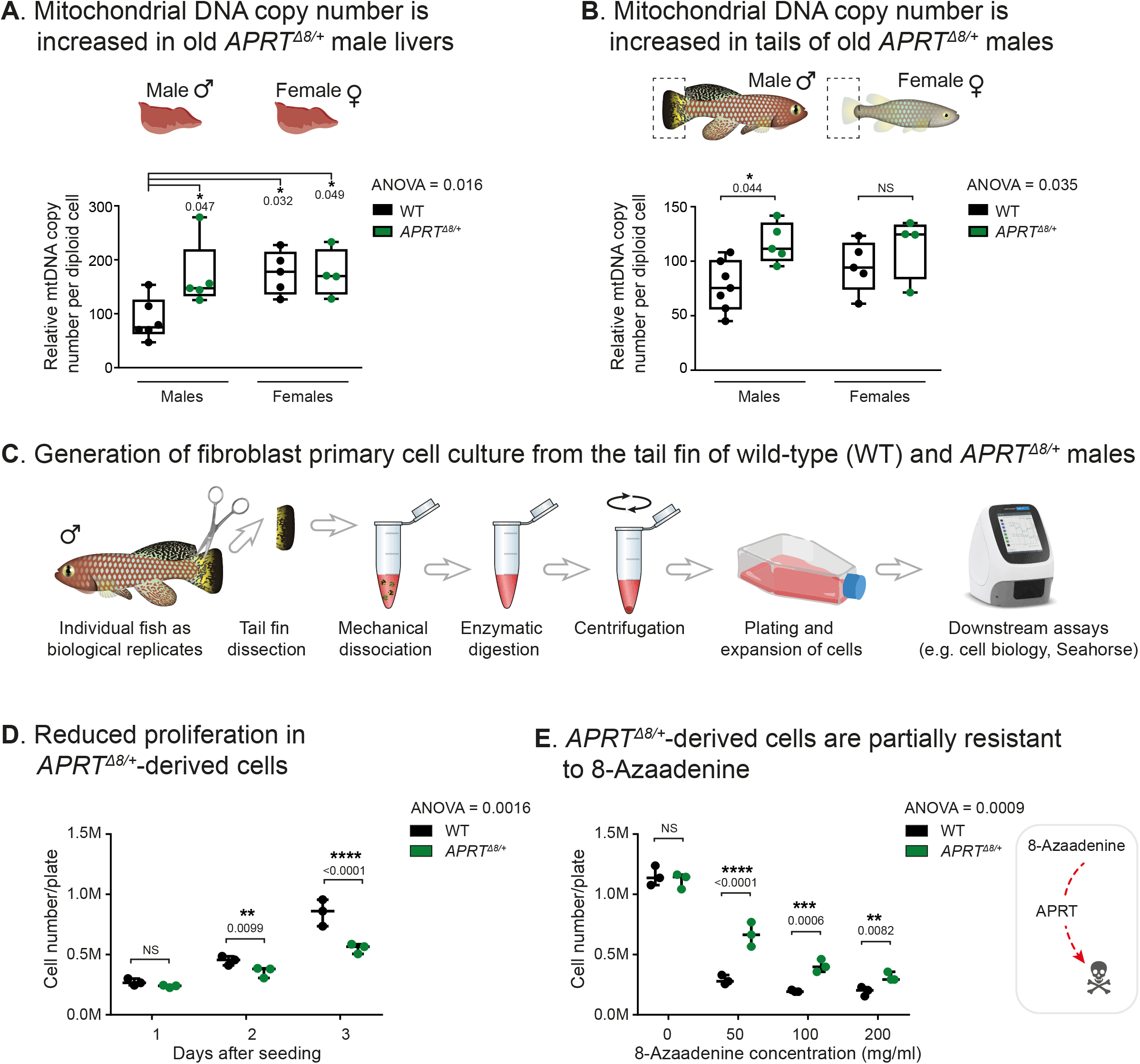
Mitochondrial DNA copy numbers are sex- and genotype-dependent. (**A-B**) Relative mitochondrial DNA (mtDNA) copy number in old fish livers (A) or tails (B) was determined by quantitative real-time PCR (qPCR) using primers for mitochondrial (*16S rRNA*) and nuclear (*CDKN2A/B*) gene. Experimental groups differ by sex (male or female) and genotype (WT or *APRT^Δ8/+^*). Each symbol represents an individual fish. Bars represent minimum and maximum, and significance was measured by one-way ANOVA and Sidak’s correction for multiple tests. Exact p-values are indicated. (**C**) Schematic illustration depicting the isolation procedure of primary fibroblasts from the tail-fin of individual fish. Isolations and downstream assays were separately performed on WT and *APRT^Δ8/+^* male fish (n=3 for each genotype). (**D-E**) Assessing cell proliferation rates (D) and resistance to 8-Azaadenine (E) in WT- and *APRT^Δ8/+^*-derived primary cells under the indicated experimental conditions. Each symbol represents a cell culture derived from an individual fish. Bars represent minimum and maximum, significance was measured by two-way ANOVA, and BKY (Benjamini, Krieger, & Yekutieli, false discovery rate) correction for multiple tests.

In search for genotype-age interactions that are shared across all samples, in males and females, highlighted four pathways involving RNA modification (Figure S5a). Such modifications, including RNA-editing (Eisenberg and Levanon, 2018), are known to be tightly linked to mitochondrial dynamics (Dhir et al., 2018; Sharma et al., 2019; Wiatrek et al., 2019). Interestingly, an in depth examination of A-to-I editing in our dataset identified an age-dependent increase in RNA-editing in males, and a decrease in females (Figure S5b-d). Although age-dependent RNA-editing was previously predicted (Porath et al., 2017), our ability to identify this phenomenon in killifish was made possible by the low polymorphic background in killifish (Valenzano et al., 2015), and by developing a new computational editing detection strategy (see Methods section).

So far, these data highlight pathways that display significant sex differences during aging, and a subset that is further altered in response to the *APRT^Δ8/+^* heterozygous mutation (particularly related to energy metabolism and mitochondrial physiology). However, evaluating whether these pathways contribute to the observed differences in lifespan, requires a cellular platform in order to explore how mitochondrial functions are affected by the *APRT^Δ8/+^* heterozygous mutation.

### *APRT^Δ8/+^* cells have reduced mitochondrial functions that can be partially rescued by adenine supplementation

To explore mitochondrial function on a cellular level, we developed primary fibroblast cultures derived from the tail fin of male WT and *APRT^Δ8/+^* fish (Figure 5c). Optimizing previously published protocols (Graf et al., 2013), allowed us to produce robust early-passage cultures, and use individual fish as biological replicates (without pooling). As expected, *APRT^Δ8/+^*-derived cells display reduced proliferation when compared to WT cells (Figure 5d) and are partially resistant to 8-azaadenine treatment (Figure 5e). This partial resistance is a consequence of the reduced ability to metabolize adenine, and therefore to produce the toxic intermediate generated by 8-azaadenine (Jones and Sargent, 1974; Sahota et al., 1987). Together, these experiments demonstrate that killifish primary cells can serve as a robust platform for cell biology.

As the next stage, we used the seahorse platform to characterize mitochondrial functions (‘Mito Stress’ assay, Figure 6a). The results indicated that mitochondrial respiration (including basal respiration and ATP production), were significantly reduced in primary cells from *APRT^Δ8/+^* fish (Figure 6b-d). Similarly, there was a significant reduction in glycolytic functions in heterozygous cells, including glycolytic capacity and glycolysis (Figure S6a-c). In contrast, mitochondrial functions, and specifically the level of ATP production, were unaltered in *AK2^Δ8/+^*-derived cells (Figure S6d). These data are in agreement with the comparable lifespan between WT and *AK2* heterozygous fish (Figure S1g,h). We next hypothesized that the observed phenotypes could be rescued by exogenously manipulating the nucleotide pools. APRT catalyzes the formation of AMP by using adenine as a precursor. Accordingly, our results revealed that adenine supplementation partially restores mitochondrial functions in heterozygous cells by increasing respiration (Figure 6c,d). These results provide support for the notion that the observed mitochondrial phenotypes are a direct consequence of APRT haploinsufficiency.

**Figure 6:**
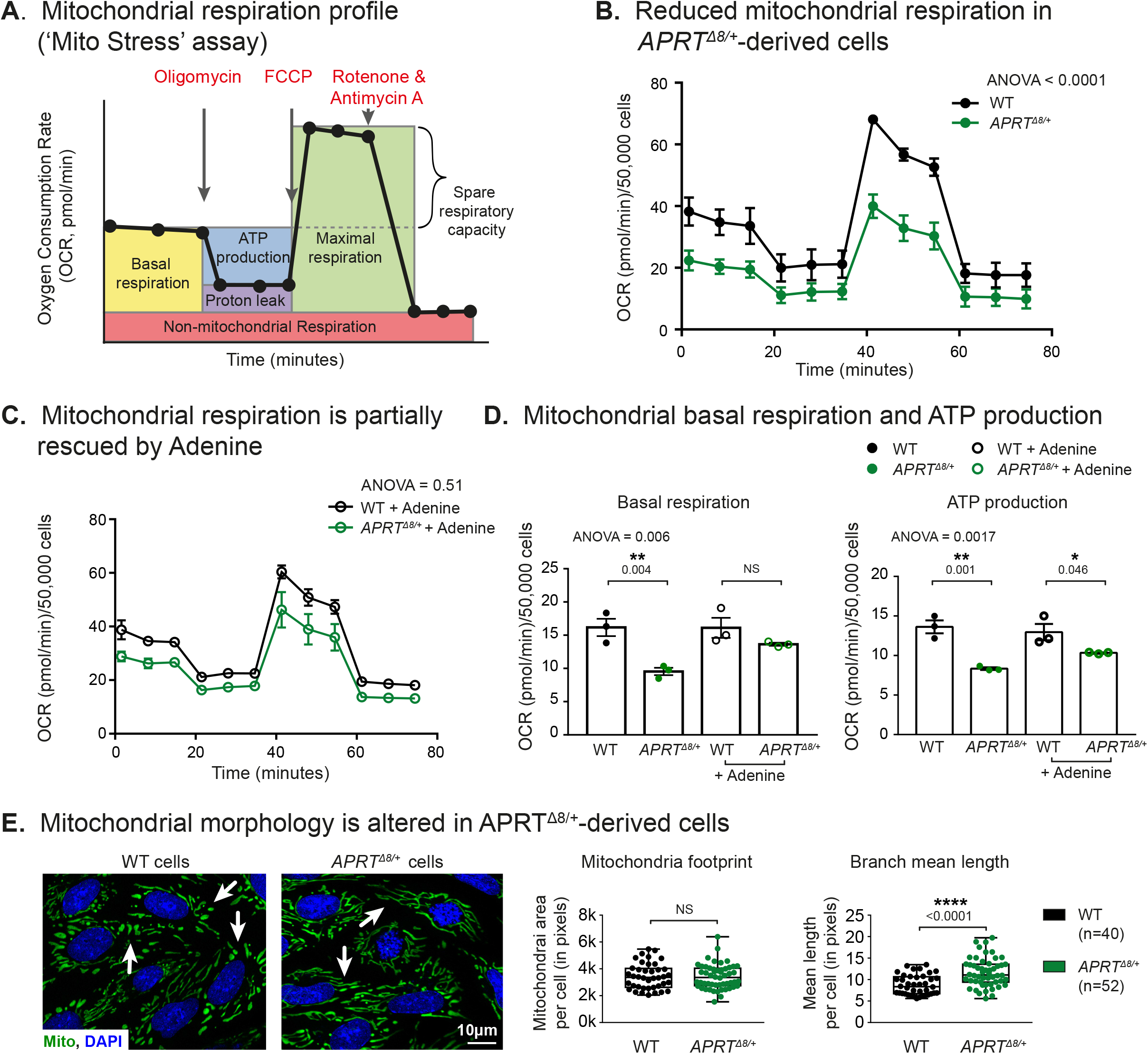
altered mitochondrial functions in *APRT^Δ8/+^* -derived cells. (**A**) Measurement of key mitochondrial parameters as a function of oxygen consumption rate (OCR, pmol/min) using the Seahorse Cell Mito Stress Test assay (Agilent). Measurements were performed under basal conditions or following the addition of the indicated inhibitors. Specific parameters, such as the rates of basal respiration and ATP production, were further normalization by actual cell numbers (D). (**B-C**) Measuring mitochondrial respiration in WT- and *APRT^Δ8/+^*-derived primary cells. Experiments were performed in control conditions (B) and adenine supplementation (C). Each symbol represents an average of three independent measurements using cells derived from three individual fish. Error bars represent mean ± SEM, and significance was measured by two-way ANOVA and BKY correction. (**D**) Basal respiration and ATP production were compared between WT- and *APRT^Δ8/+^*-derived primary cells for control (B) and adenine supplementation (C). Each symbol represents an individual fish. Error bars represent mean ± SEM, and significance was measured one-way ANOVA and Tukey post-hoc. (**E**) Left: live imaging of WT- and *APRT^Δ8/+^*-derived primary cells using mitochondrial staining (MitoView, green), and Hoechst (blue). Scale bar: 10 µm. Right: quantification of mitochondrial footprint (volume) and branch mean length was performed using MiNA macro plugin (Fiji - ImageJ). Data were obtained from 3 independent animals per genotype, with the indicated number of individualized mitochondria. Error bars represent mean ± SEM and the significance were measured by an unpaired t-test.

Could the observed phenotypes be attributed simply to changes in mitochondrial numbers, as seen *in-vivo* (Figures 5a)? To test this, we investigated mitochondrial morphology in the cultured cells. We observed that although mitochondrial branch length is slightly increased in heterozygous cells, mitochondrial footprint remains essentially the same regardless of genotype (Figure 6e). These findings were further supported by direct measurement of mtDNA copy numbers in cells, which revealed comparable counts in WT and heterozygous cells (Figure S6e). Surprisingly, mitochondrial copy number measurements were sex-independent in cultured cells, demonstrating similar results in male- and female-derived cells (Figure S6e). Similarly, characterizing mitochondrial respiration in female-derived cells revealed that the APRT-dependent effect was also comparable between males and females (Figure S6f). Thus, suggesting that the observed sex-biased phenotypes in mitochondrial copy numbers could be mediated by cell non-autonomous mechanism, such as sex hormones (Mohapatra et al., 2020).

To investigate how aging affects the cellular response to circulating sex hormones we used gene set variation analysis (GSVA, Figure S6g) (Hänzelmann et al., 2013). Interestingly, although steroid biosynthesis (the precursor for gonadal sex hormones), was significantly downregulated in old heterozygous males (when compared to old wild-type males), the cellular response to estrogen was upregulated (Figures 4b, S6g). And indeed, short-term exposure of wild type fish to estradiol (E2), significantly increased mitochondrial copy number, specifically in males (Figure S6h).

Together, our findings suggest that the observed mitochondrial copy number phenotypes (which are genotype- and sex-dependent), could be linked to circulating *in-vivo* signals, such as sex hormones. In contrast, in heterozygous cultured cells, mitochondrial functions are inhibited cell autonomously in a sex-independent manner. As pharmacological or genetic inhibition of mitochondrial functions is a conserved longevity intervention (Baumgart et al., 2016; Lee et al., 2003; Sun et al., 2016; Tavallaie et al., 2020; Wang and Hekimi, 2015), our findings could provide a possible pro-longevity mechanism downstream to APRT. Interestingly, our data predicts an inhibition of the mitochondrial respiratory chain, specifically in old *APRT^Δ8/+^* heterozygous males (Figure 4b). However, the question remains as to which physiological mechanisms are directly involved in the male-specific increased longevity.

### Increased AMPK activity and altered nucleotide ratios in *APRT* mutant cells produces a fasting-like state

The data so far suggest that *APRT^Δ8/+^*-derived cells experience altered energy metabolism. To further explore whether this condition resembles fasting on the protein level, we validated a panel of specific antibodies directed against key members of the vertebrate fasting response (Figure S7a,b). These were used to assess the phosphorylation of classical AMPK targets in WT and *APRT^Δ8/+^*-derived cells, under either basal conditions, or low serum (serum starvation, Figure 7a). Our results indicate that under basal conditions, *APRT^Δ8/+^*-derived cells display a significant activation of the AMPK pathway (either direct or indirect), which resembles the activation observed in WT cells under serum starvation. This increase is further heightened under serum starvation conditions, suggesting that the *APRT^Δ8/+^* mutation can sensitize cells to nutrient levels and induce a fasting-like state (even under basal conditions).

**Figure 7:**
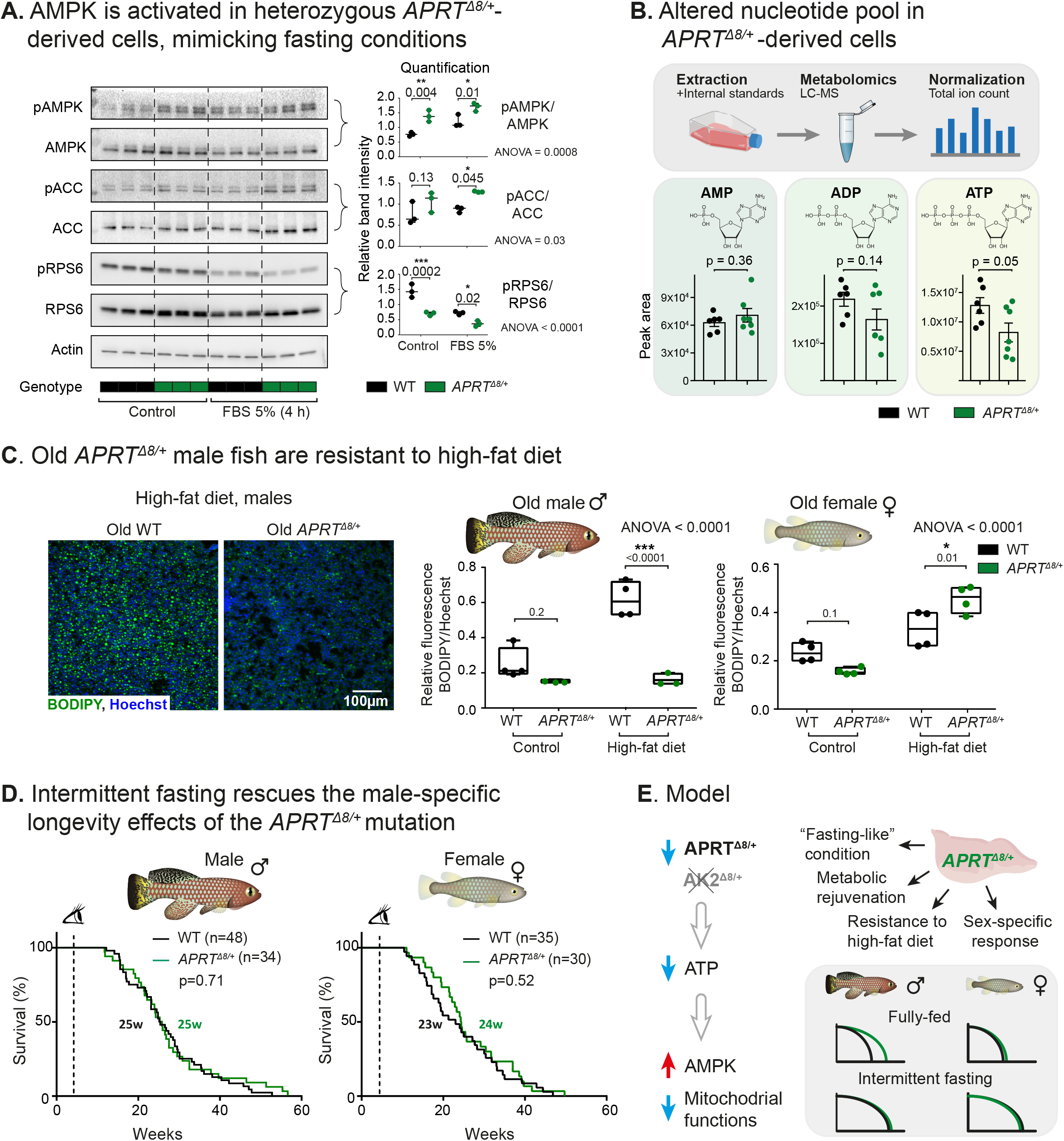
A sex-specific fasting-like state mediates the longevity effect of the APRT mutation. (**A**) Western blot (left) of primary metabolic pathways in WT- and *APRT^Δ8/+^*-derived primary cells. Quantification (Fiji – ImageJ, right) represents the ratio between the phosphorylated and non-phosphorylated forms. Experiments were performed under either control or fasting (5% FBS, 4 h) conditions. Data from 3 biological replicates for each experimental condition. Bars represent minimum and maximum, significance was measured by one-way ANOVA and Sidak’s correction for multiple tests. Exact p-values are indicated. AMPK: AMPKα; pAMPK: Phospho-AMPKα (Thr172); ACC: Acetyl-CoA Carboxylase; pACC: Phospho-Acetyl-CoA Carboxylase (Ser79); RPS6: S6 Ribosomal Protein; pRPS6: Phospho-S6 Ribosomal Protein (Ser235/236). (**B**) Direct measurement of the nucleotide pool (relative concentration of AMP, ADP, and ATP) in WT- and *APRT^Δ8/+^*-derived primary cells using liquid chromatography–mass spectrometry (LC–MS). Data were obtained from 6-7 biological replicates for each experimental condition, and normalized by total ion count (TIC). Bars represent mean ± SEM, unpaired t-test and exact p-values are indicated. (**C**) Hepatic lipid accumulation and quantification in response to high-fat diet. Left: BODIPY staining (green) of liver section. Nuclei were stained by Hoechst (blue). Scale bar represents 100 µm. Right: Quantification of the relative fluorescence intensity of BODIPY/Hoechst (Fiji-ImageJ). Experimental groups differ by sex (males or females), feeding condition (control or high-fat diet), and genotype (WT or *APRT^Δ8/+^*). n=3-4 for each experimental condition. Bars represent minimum and maximum, and significance was measured by one-way ANOVA and Sidak’s correction for multiple tests. Exact p-values are indicated. (**D**) Lifespan of WT (black) and *APRT^Δ8/+^* (green) fish performed separately for males (left) and females (right) under intermittent fasting conditions. pvalues for differential survival in log-rank tests, median survival, and fish numbers are presented. See Table S1 for additional information. (**G**) A model suggesting that APRT, a member of the AMP biosynthesis pathway, could function as a regulator for vertebrate longevity. Mutating *APRT* induces a sex-specific longevity effect, mediated by restoring metabolic homeostasis, modulating mitochondrial functions, and inducing a fasting-like state that involves a distinct AMPK signature.

In fish, a genotype-dependent response to fasting was also observed on the protein level, which was evaluated by phosphorylation levels of ribosomal protein S6 (pRPS6, a readout for mTOR signaling, Figure S7c), as well as by specific transcriptional signatures associated with the mTOR and AMPK pathways (Figure S7d, left). Examining the expression levels of individual AMPK complex subunits (the α, β, and γ subunits, and their respective isoforms), revealed a distinct genotype-dependent compositional change. Specifically, only a single isoform, PRKAG2 (protein kinase AMP-activated non-catalytic subunit γ2) displayed an over 2-fold increase between the livers of *APRT^Δ8/+^* and WT fish (p values = 0.03 and 0.07, for fully-fed young and old fish, respectively. Figure S7d, right). These findings propose that under low-energy conditions (i.e. the fasting-like state of heterozygous male fish), PRKAG2 might be involved in promoting metabolic homeostasis.

Mutating *APRT*, which is part of the adenosine nucleotide synthesis pathway, might be expected to alter the ratios of AMP, ADP, and ATP. Furthermore, data in invertebrate models suggests that the ratios of these products (e.g., AMP:ATP) are altered in response to dietary restriction, and that such alterations could be predictive of the organismal lifespan (Apfeld et al., 2004; Greer et al., 2007; Stenesen et al., 2013). It was therefore of interest to explore whether this effect is also conserved in vertebrates. For this purpose, we analyzed the primary cultures by liquid chromatography mass spectrometry (LC–MS). The results revealed that ATP levels are significantly decreased in *APRT^Δ8/+^*-derived cells (p = 0.05, t-test), while there was a non-significant trend to lower ADP ratios (p = 0.14, t-test), and the AMP levels seem to be unchanged (Figures 7b). Overall, these data imply that the *APRT^Δ8/+^*-derived cells exist in a low energy state. We were next curious to test whether a similar fasting-like state also occurs in the male *APRT^Δ8/+^* heterozygous fish.

### A sex-specific fasting-like state is observed in male *APRT^Δ8/+^* fish

A variety of physical traits have been linked to longevity interventions that exert their effect by manipulating organismal metabolism (Fontana et al., 2010; Lopez-Otin et al., 2013; Smith et al., 2020). These include reduced body size and fecundity, delayed maturity, and resistance to a high-fat diet. Although *APRT^Δ8/+^* mating pairs exhibited no significant differences in fecundity (Figure S1b), male heterozygous fish were slightly smaller than their WT counterparts (Figure S7e) and experienced delayed maturity (Figure S7f). In contrast, the growth of female fish was unaffected by the *APRT ^Δ8/+^* mutation (Figure S7e). As the next step, we developed a short-term high-fat diet regimen, which efficiently increases the amount of hepatic lipid droplets, specifically in old fish (Figures S7g,h). Applying this protocol to both male and female fish revealed that in contrast to females, old *APRT^Δ8/+^* males are completely resistant to high-fat diet, and maintained hepatic lipid levels comparable to those of untreated control fish (Figures 7c, and Table S1 for a summary of pathological findings).

### Intermittent fasting eliminates the longevity benefits mediated by the *APRT* mutation

Our data demonstrate that the *APRT^Δ8/+^* mutation induces a fasting-like physiological state, specifically in male fish. This has led us to hypothesize that the *APRT^Δ8/+^*-mediated longevity benefits might be effective only under normal feeding conditions, and may be negated by fasting. Excitingly, although the median lifespan of both male and female fish was significantly extended following life-long intermittent fasting (by ∼70%, in both male and female WT fish, Figure S7i), we could not detect any further extension in *APRT^Δ8/+^*male fish (log-rank test, p = 0.71, Figure 7d). This observation suggests that the longevity effect achieved by intermittent fasting were dominant over the APRT-dependent sex differences. It is important to note that the effects of dietary restriction themselves are thought to be partially mediated by sex-specific molecular responses (Figures 4d, S4c) (Kane et al., 2018) (see **Discussion**).

Together, our findings demonstrate that mutating *APRT*, a member of the AMP biosynthesis pathway, induces a sex-specific longevity effect. This effect is mediated by restoring metabolic homeostasis, modulating mitochondrial functions, and by inducing a fasting-like state that involves distinct AMPK downstream regulators. In conclusion, our data suggest that the AMP/AMPK axis, through the AMP salvage pathways, could function as a rheostat for vertebrate healthspan and longevity (Figure 7e).

## Discussion

Sexual dimorphism has been detected in many metabolic pathways. These sex differences are evolutionarily linked to the unique metabolic requirements and energy partitioning required during reproductive life (Austad, 2011; Mauvais-Jarvis, 2015; Naqvi et al., 2019). With age, these characteristics might also contribute to the sex-biased predisposition to various diseases, such as Alzheimer’s disease and type 2 diabetes (Hägg and Jylhävä, 2021; Tramunt et al., 2020). Accordingly, genetic manipulation of several longevity pathways in mice confers sex-specific longevity responses (Ashpole et al., 2017; Baar et al., 2016; Green et al., 2021; Kanfi et al., 2012; Sharples et al., 2015; Sun et al., 2017).

Similarly, pharmacological longevity interventions, such as via metformin administration (a non-specific AMPK activator (Pernicova and Korbonits, 2014)), produces trending male-specific longevity effects (https://phenome.jax.org/projects/ITP1). Exploring sex differences in longevity is feasible in other vertebrate models, including dogs (Bray et al., 2021; Creevy et al., 2022), and could be further examined in human clinical trials (Barzilai, 2017; Kulkarni et al., 2020). Thus, better understanding the molecular mechanisms behind sex-specific longevity differences could provide unique opportunities for manipulating health and disease.

By exploring the basis for the sex-dependent longevity of the *APRT* mutants, we identified mitochondrial functions as a possible molecular mechanism. Accordingly, mitochondrial DNA copy numbers are increased, specifically in old *APRT* heterozygous males. Indeed, mitochondria function and copy number are known to exhibit an evolutionarily-conserved sex bias (Ballard et al., 2007; Borras et al., 2007; Di Florio et al., 2020; Justo et al., 2005; Klinge, 2008; Kristensen et al., 2019; Valle et al., 2007; Ventura-Clapier et al., 2017). It is possible that in females, mitochondrial and numbers are naturally optimized (Figures 5a) (potentially due to the high metabolic demand of egg production). Thus, an increase in mitochondria numbers by the *APRT* mutation, is more likely to be apparent in males.

Insight into how metabolic signals are differently integrated during aging in males and females opens up new possibilities for the design of pro-longevity interventions. However, a mechanistic understanding of how these signals are transmitted between tissues, cells, and organelles is still lacking. Our findings uncouple between the signals that regulate mitochondrial function to the ones that promote mitochondrial biogenesis. Specifically, the mitochondrial functions observed here are regulated in a cell-autonomous manner, and are genotype dependent (regardless of whether the cells concerned were isolated from male or female fish, Figures 6, S6, S7).

In contrast, mitochondrial DNA copy numbers appear to be dependent on physiological signals, as the *in-vivo* phenotypes were not present in primary cells. Recent studies suggest sex hormones as a possible mediator (Besse-Patin et al., 2017; Hamilton et al., 2016; Zhou et al., 2020), and accordingly, estrogen exposure alone was sufficient to replicate our findings (Figure S6h). Further exploring these signals in future studies could provide a powerful tool for systemic alteration of mitochondrial biogenesis *in-vivo*.

The longevity effects of intermittent fasting were dominant over the APRT-dependent sex differences, although this must be reconciled with the observation that intermittent fasting affects both sexes, while APRT has an impact only on male fish. A possible explanation, as implicated by our findings (Figures 4d and S4c), and by increasing evidence from rodents, is that the beneficial effects of dietary restriction are partially mediated by sex-specific downstream regulators (Green et al., 2021; Kane et al., 2018). Notably, in genetic mouse models of APRT deficiency, males display increased susceptibility (Evan et al., 2001), suggesting that the sex differences in AMP metabolism could be evolutionarily conserved (Naqvi et al., 2019).

Our findings propose the APRT enzyme as a promising pharmacological target for mimicking low energy conditions, for replicating the benefits of fasting, and for promoting vertebrate longevity. However, although significant efforts have been invested in *Leishmania* research (Boitz et al., 2012), a selective APRT inhibitor is still missing. Intriguingly, APRT inhibition by the APRT heterozygous mutation is predicted to actually reduce AMP levels and therefore *decrease* AMP:ATP ratios. Thus, the observed AMPK activation is counterintuitive under steady-state conditions, as seen in *APRT* heterozygous *Drosophila* (Camici et al., 2018; Stenesen et al., 2013). However, this phenomenon could simply suggest that under low AMP production rates, the cell is more sensitive to spikes in energy consumption (e.g. during cell division), or there is an as yet unidentified regulatory mechanism for AMPK (Yee et al., 2014). Interestingly, under normal energy conditions, an additional AMPK-related mechanism is required for restoring metabolic homeostasis during vertebrate aging (see accompanying paper by Ripa et al).

Using the turquoise killifish model, we identify that AMP biosynthesis is a sex-specific regulator of metabolic health and vertebrate lifespan. However, to what extent are our findings evolutionarily conserved? A recent study (Lin et al., 2021) identified longevity-associated rare coding variants that converged on a number of pathways, including the AMPK signaling pathway. Together, our findings provide a global understanding into how the AMP/AMPK axis function as a regulator of vertebrate metabolism and longevity.

## Acknowledgements

We thank the Harel lab, Sagiv Shifman, Eran Meshorer, Anne Brunet, and Berenice Benayoun for stimulating discussion and feedback on the manuscript. We thank Ella Yanay and Ashayma Abu-tair for help with killifish maintenance, Yaakov Nahmias and Konstantinos Ioannidis for help with the Seahorse assay, Naomi Melamed-Book and Rachel Rosen from the core facilities (HUJI), Ifat Abramovich, Eyal Gottlieb, Sergey Malitsky and Maxim Itkin for help with mass spectrometry, and Param Priya Singh and Anne Brunet for advice with the GSEA code. Supported by the Zuckerman Program (I.H.), Abisch-Frenkel Foundation 19/HU04 (I.H.), ISF 2178/19 (I.H.), Israel Ministry of Science 3-17631 (I.H.), 3-16872 (I.H.), the Moore Foundation GBMF9341 (I.H.), BSF-NSF 2020611 (I.H.), the Israel Ministry of Agriculture 12-16-0010 (I.H), and the Lady Davis Postdoctoral Fellowship (G.A).

## Author Contributions

G.A. and I.H. designed the study. G.A. performed experiments with help from U.G, A.O.G., T.A., and T.L.. T.A. designed and performed the analysis of multi-omic datasets and performed statistical analysis with help for G.A. K.S. performed the RNA editing predictions under the supervision of E.Y.L. A.V. performed injections and generated the AK2 and APRT mutant killifish lines. M.S. assisted G.A with the histology, and J.D. advised regarding GWAS predictions. T.A., G.A., and I.H. wrote the manuscript. All authors commented on the manuscript.

## METHODS

Detailed methods are provided in the online version of this paper and include the following:

## KEY RESOURCES TABLE

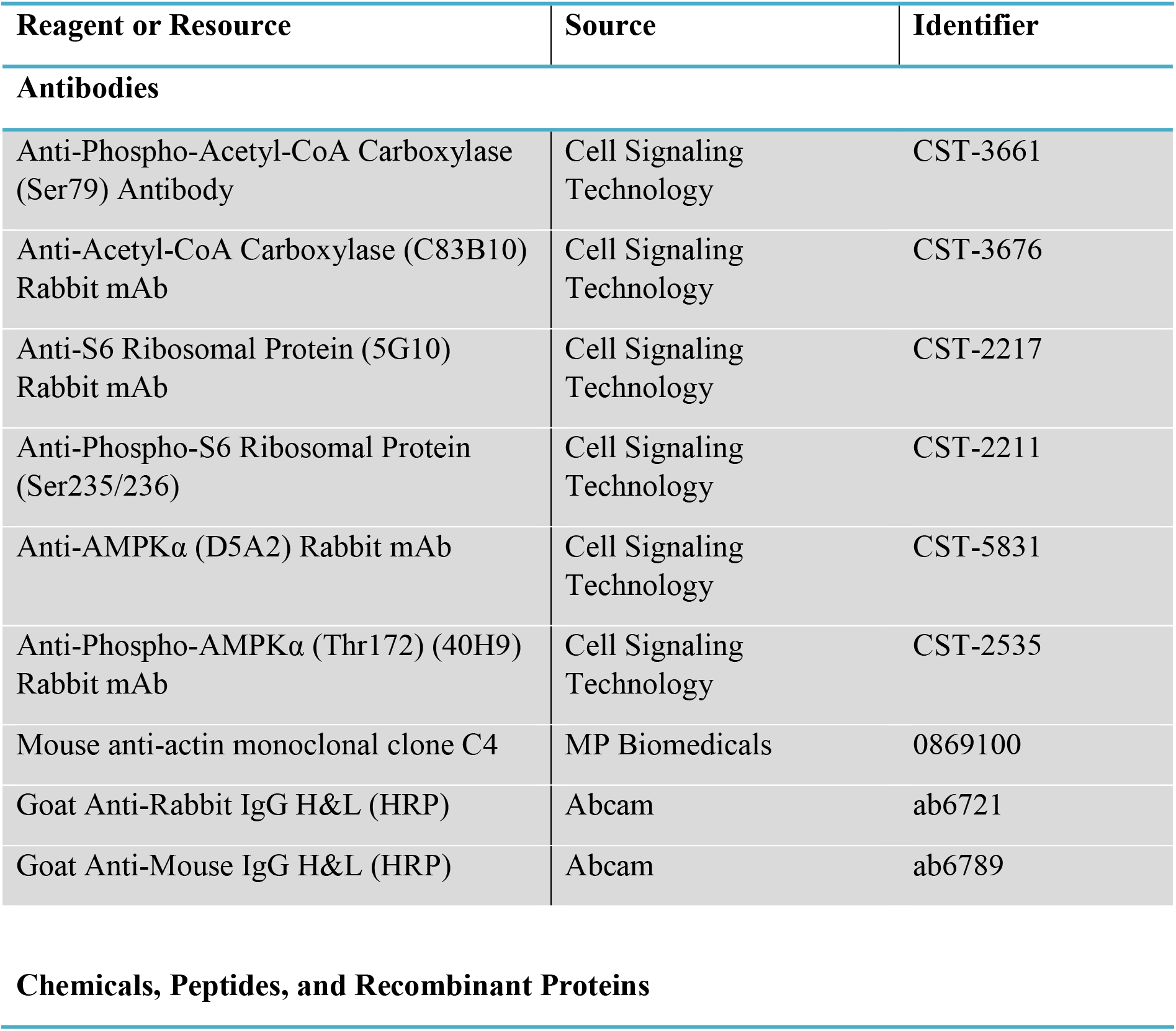
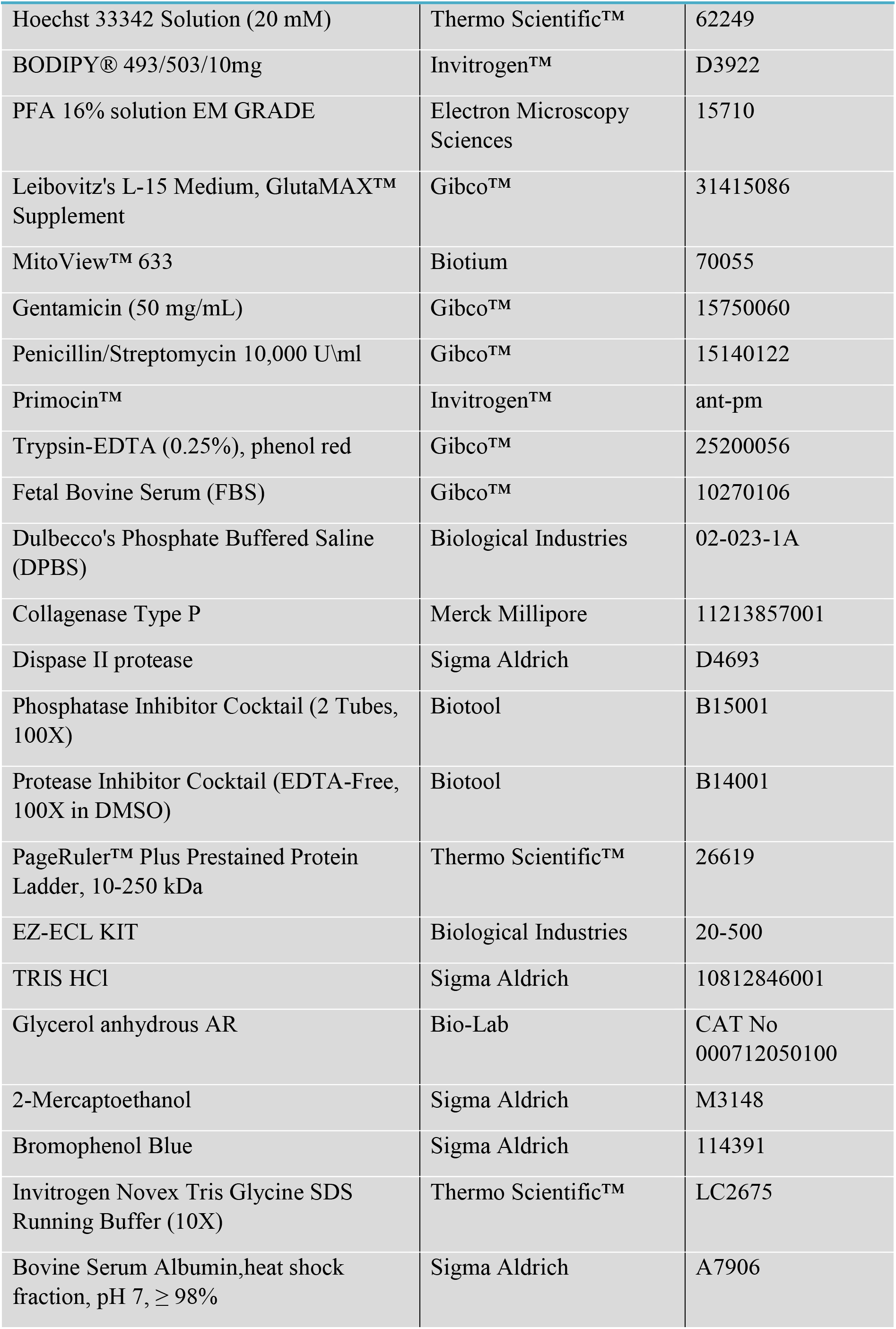
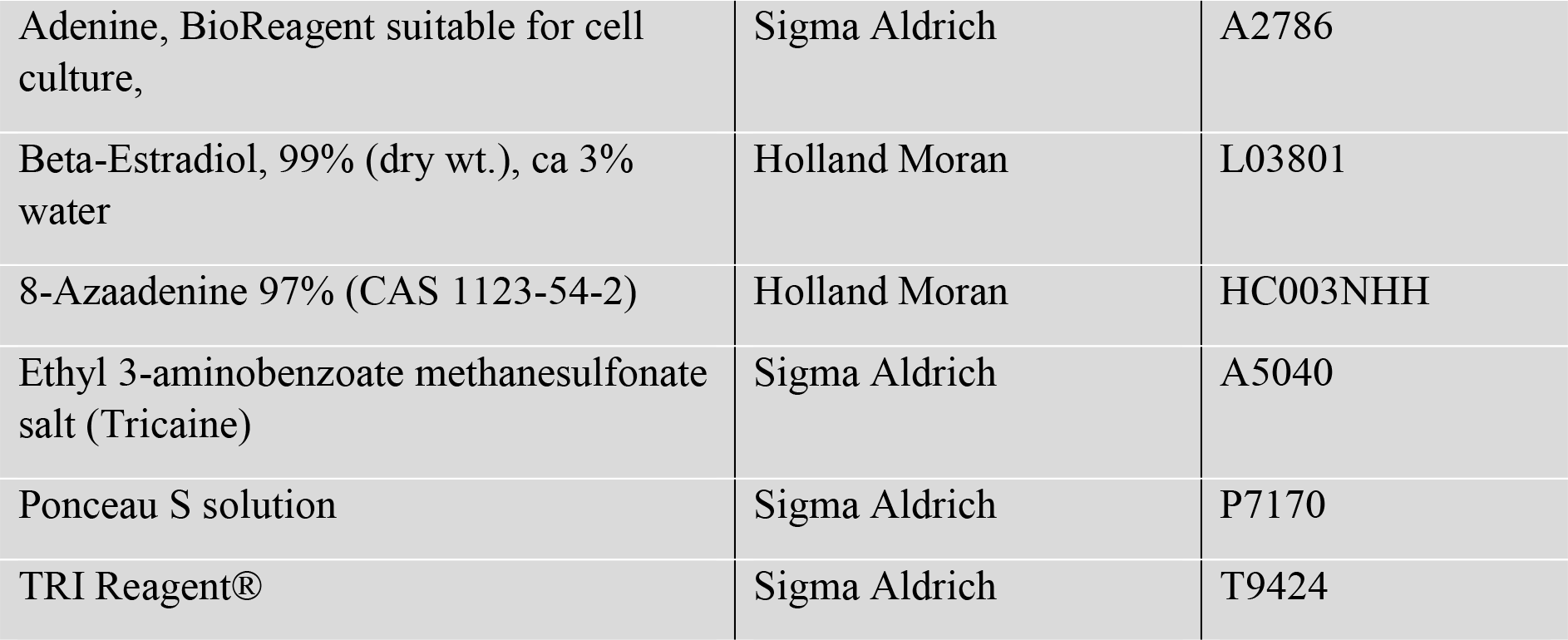

### Software and Algorithms

|  |  |  |
| --- | --- | --- |
| GraphPad Prism (version 7 for Windows) | GraphPad software | https://www.graphpad.com/ |
| Fiji 1.53c (plus) | NIH | https://imagej.net/Fiji |
| Matlab R2021a | The MathWorks, Inc. | https://www.mathworks.com/ |
| FlowView confocal software | Olympus, Japan |  |
| Olympus cellSens Dimension 3.1 | Olympus |  |
| FastQC | (Andrews, 2017) | Version 0.11.8 |
| fastx-toolkits |  | Version 0.0.13 |
| Trim Galore! | (Krueger, 2015) | Version 0.6.4 |
| Cutadapt | (Martin, 2011) | Version 3.4 |
| STAR | (Dobin et al., 2013) | Version 2.7.6a |
| edgeR | (McCarthy et al., 2012; Robinson et al., 2010) | Version 3.32.1 |
| clusterProfiler | (Yu et al., 2012) | Version 3.18.1 |
| GSVA | (Hänzelmann et al., 2013) | Version 1.40.1 |
| Trimmomatic | (Bolger et al., 2014) | Version 0.39 |
| RepeatMasker | (Chen et al., 2004) | Version 4.1.0 |
| EDTA | (Ou et al., 2019) | Version 1.9.6 |
| RNA Editing Indexer | https://github.com/a2iEditing/RNAEditingIndexer |  |
| Salmon | (Patro et al., 2017) | Version 1.4 |
| BWA | (Li and Durbin, 2009) | Version 0.7.17 |

### Critical Commercial Assays

|  |  |  |
| --- | --- | --- |
| Seahorse XFp FluxPak | Agilent Technologies | 103022-100 |
| Seahorse XFp Cell Mito Stress Kit | Agilent Technologies | 103010-100 |
| Seahorse XF Glycolysis Stress Kit | Agilent Technologies | 103017-100 |
| Novex™ WedgeWell™ 4 to 20%, Tris-Glycine, 1.0 mm, Mini Protein Gel, 15-well | Invitrogen™ | XP04205BOX |
| Verso cDNA Synthesis Kit | Thermo Scientific™ | AB1453 |
| Pierce™ BCA Protein Assay Kit | Thermo Scientific™ | 23225 |
| Fast SYBR™ Green Master Mix | Applied Biosystems™ | 4385612 |
| Direct-Zol RNA Miniprep | Zymo | ZR-R2050 |
| QIAamp DNA Micro Kit | Qiagen | 56304 |
| Kapa RNA HyperPrep | ROCHE | 08105936001 |
| NextSeq 500/550 High Output Kit v2.5 (75 Cycles) | Illumina | 20024906 |

### Oligonucleotides

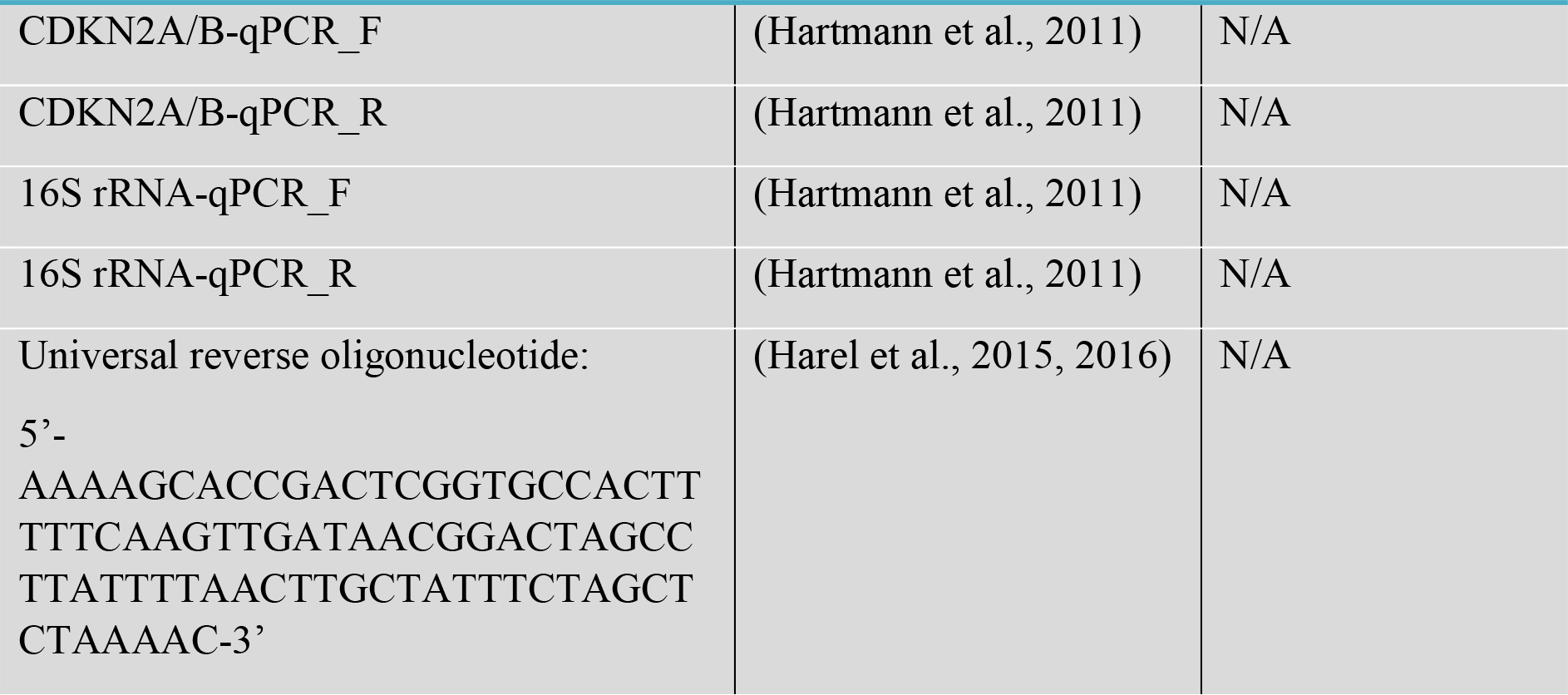
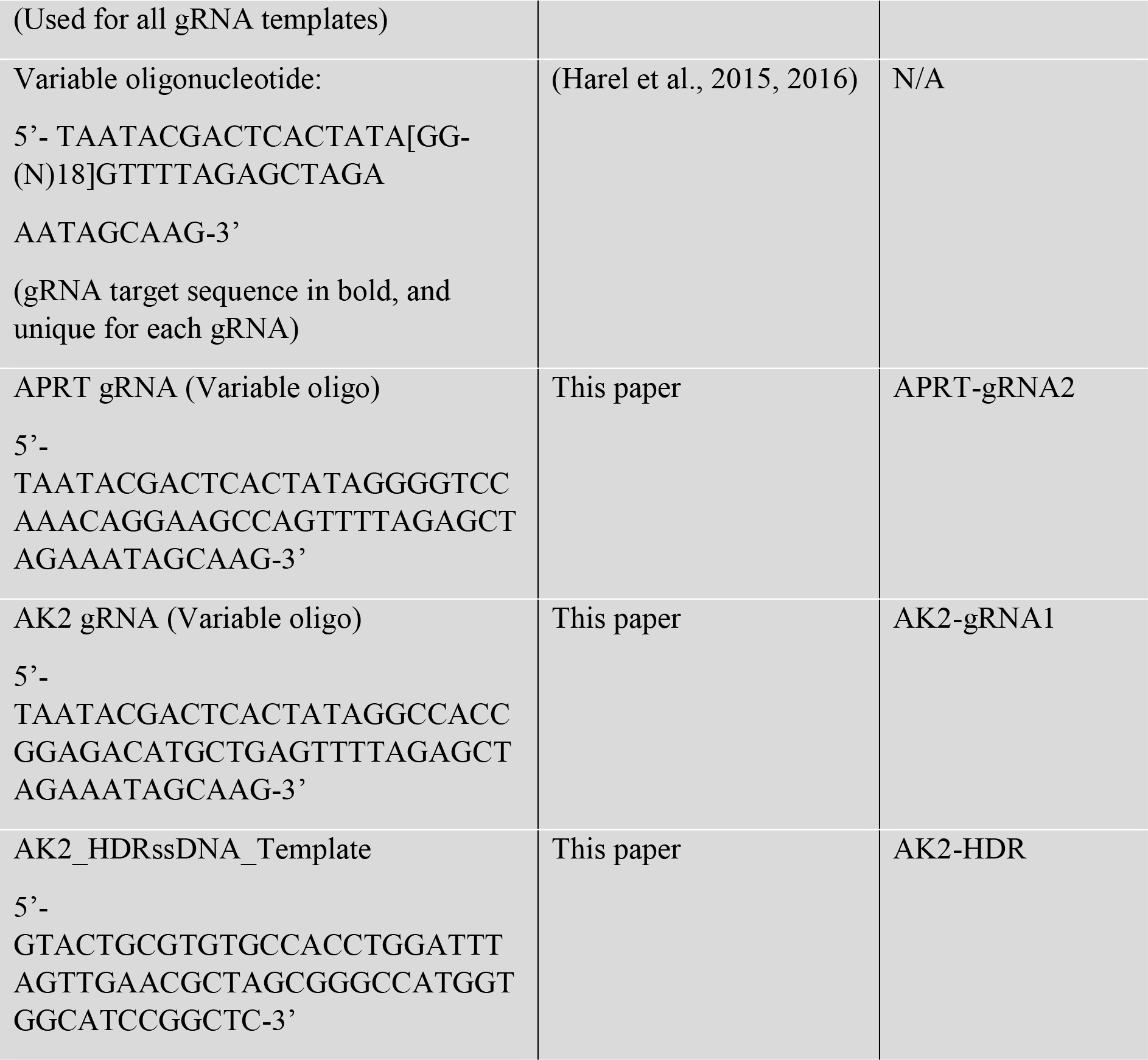

### Fish

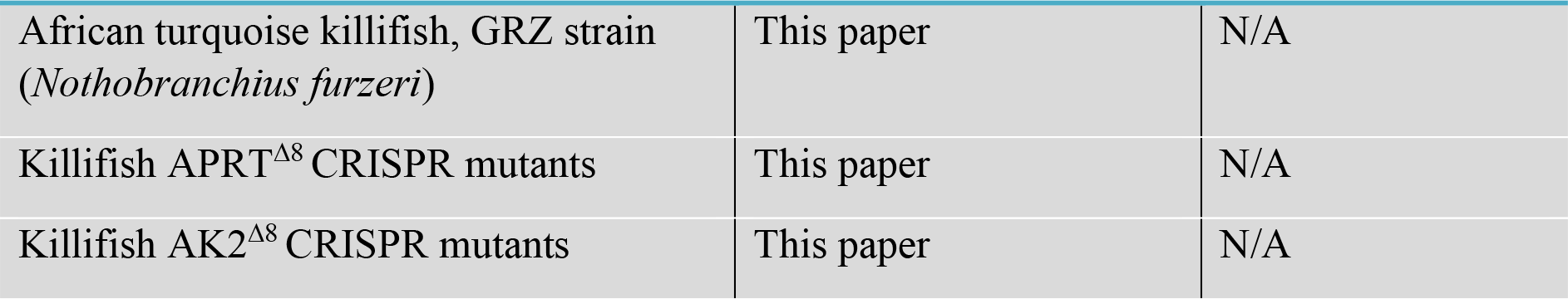

|  |  |  |
| --- | --- | --- |
| African turquoise killifish, GRZ strain<br>(*Nothobranchius furzeri*) | This paper | N/A |
| Killifish APRT<sup>Δ8</sup> CRISPR mutants | This paper | N/A |
| Killifish AK2<sup>Δ8</sup> CRISPR mutants | This paper | N/A |

## RESOURCE AVAILABILITY

### Lead Contact

Further information and requests for resources and reagents should be directed to and will be fulfilled by the Lead Contact, Itamar Harel.

### Data and Code availability

All raw RNA sequencing data, as well as processed datasets could be found in the GEO database, accession number GSE190757. All Lipidomic and metabolomic datasets could be found in Supplementary Table S3. The codes and results supporting the current study are available in the GitHub repository for this paper https://github.com/Harel-lab/APRT-sex-differences.

## EXPERIMENTAL MODEL AND SUBJECT DETAILS

### African turquoise killifish strain, husbandry, and maintenance

The African turquoise killifish (GRZ strain) were housed as previously described (Astre et al., 2022; Harel et al., 2016). Fish were housed at 28°C in a central filtration recirculating system with a 12 h light/dark cycle at the Hebrew University of Jerusalem (Aquazone ltd, Israel). Until the age of 2 weeks, fish were exclusively fed with live Artemia (#109448, Primo). Starting week 3, fish were fed three times a day on weekdays (and once a day on weekends), with GEMMA Micro 300 Fish Diet (Skretting Zebrafish, USA), supplemented with Artemia twice a day. In these conditions, killifish lifespan was approximately 4-6 months. Both the APRT and AK2 loss-of-function alleles were maintained as heterozygous and propagated by crossing with wild-type fish. All turquoise killifish care and uses were approved by the Subcommittee on Research Animal Care at the Hebrew University of Jerusalem (IACUC protocol #NS-18-15397-2).

### CRISPR/Cas9 target prediction and gRNA synthesis

CRISPR/Cas9 genome-editing protocols were performed according to (Astre et al., 2022). In brief, for targeting APRT and AK2, conserved regions that were upstream of functional or active protein domains were selected. gRNA target sites were identified using CHOPCHOP (https://chopchop.rc.fas.harvard.edu/) (Labun et al., 2019), and were as follows (PAM sites are in bold): APRT Exon 3: 5’-GGGGTCCAAACAGGAAGCCA**CGG**-3’; AK2 Exon 2: 5’-GGCCACCGGAGACATGCTGA**GGG**-3’. Design and hybridization of variable oligonucleotides (which are gRNA-specific) with a universal reverse oligonucleotide was performed according to (Astre et al., 2022), and the resulting products were used as a template for *in vitro* transcription. gRNAs were *in vitro* transcribed and purified using the MAXIscript T7 kit (ThermoFisher # AM1312), according to the manufacturer’s protocol.

### Production of Cas9 mRNA

Experiments were performed according to (Astre et al., 2022). The pCS2-nCas9n expression vector was used to produce Cas9 mRNA (Addgene, #47929) (Jao et al., 2013). Capped and polyadenylated Cas9 mRNA was *in vitro* transcribed and purified using the mMESSAGE mMACHINE SP6 ULTRA (ThermoFisher # AM1340).

### Single-Stranded DNA Template for Homology-Directed Repair (HDR)

Homology-directed repair (HDR) experiments for the *AK2* gene, a ssDNA template was designed to contain short homology arms (∼20 bp) surrounding the gRNA target, and included a novel sequence that replaced the endogenous gRNA target with an NheI restriction site: 5’-GTACTGCGTGTGCCACCTGGATTTAGTTGAACGCTAGCGGGCCATGGTGGCATCC GGCTC-3’. The ssDNA template was commercially synthesized and purified prior to injection (QIAquick Nucleotide Removal Kit, QIAGEN) according to (Astre et al., 2022).

### Microinjection of turquoise killifish embryos and generation of mutant fish using CRISPR/Cas9

Microinjection of turquoise killifish embryos was performed according to (Astre et al., 2022). Briefly, nCas9n-encoding mRNA (300 ng/μL) and gRNA (30 ng/μL) were mixed with phenol-red (P0290, Sigma-Aldrich) and co-injected into one-cell stage fish embryos. Sanger DNA sequencing was used for detecting successful germline transmission on F1 embryos. The genomic area encompassing the targeted site (∼600 bp) was PCR-amplified using the following primer sequences: APRT_F: 5’-TTCCCTCTTTACTGACGTCTCA-3’; APRT_R: 5’-GAAAAATTCCCACAGTAAGAATGAA-3’ and AK2_F: 5’-CCCAGGTTCTCTGTTGCATT-3’; AK2_R: 5’-GCGGTTTCCACACAAGAACT-3’. Fish with desired alleles were maintained as stable lines and further outcrossed to minimize potential off-target effects.

## METHOD DETAILS

### Organ isolation

Individual killifish, according to the specified age, gender, genotype, and feeding condition, were euthanized in 400 mg/L of Tricaine in system water. Animals were dissected on ice under a binocular stereo microscope (Leica S9E) according to (Astre et al., 2022). Whole livers were harvested, cut in half, and placed in two separate tubes. For each fish, one tube was processed for metabolomic and lipidomic profiling, while the other was processed for RNA sequencing (see below). Following weight measurements, tubes were snap-frozen in liquid nitrogen, and stored in −80°C until all samples were collected. All samples were collected during the morning time, between 9 am to 12 pm, to reduce the potential confounding effects driven by circadian rhythms.

### Lipid and polar metabolic profiling

#### Sample Preparation

Extraction and analysis of lipids and polar metabolites was performed at the Life Sciences Core Facilities, Metabolic Profiling Unit (Weizmann Institute of Science), as previously described in (Malitsky et al., 2016; Salem et al., 2017) with some modifications. Briefly, liver samples were lyophilized, ground to powder and mixed with 1 mL of a pre-cooled (−20°C) homogenous methanol:methyl-tert-butyl-ether (MTBE) 1:3 (v/v) mixture, containing the following internal standards: 0.1 μg/mL of Phosphatidylcholine (17:0/17:0) (Avanti), 0.4 μg/mL of Phosphatidylethanolamine (17:0/17:0, 0.15 nmol/mL of Ceramide/Sphingoid Internal Standard Mixture I (Avanti, LM6005), 0.0267 µg/mL d5-TG Internal Standard Mixture I (Avanti, LM6000) and 0.1 μg/mL Palmitic acid-13C (Sigma, 605573). The tubes were vortexed and then sonicated for 30 min in ice-cold Transsonic 460/H sonication bath (Elma) at 35 kHz (taken for a brief vortex every 10 min). Then, UPLC-grade water: methanol (3:1, v/v) solution (0.5 mL), containing internal polar metabolite standards (C13 and N15 labeled amino acids standard mix, Sigma, 767964) were added to the tubes. Following 5 min centrifugation at maximum speed, the upper organic phase was transferred into 2 mL Eppendorf tube. The polar phase was re-extracted as described above, with 0.5 mL of MTBE. Both parts of organic phase were combined and dried (at 21 °C setting) using a Refrigerated CentriVap Concentrator (Labconco) and then stored at −80°C until analysis. Lower, polar phase, was similarly lyophilized and stored at −80°C until analysis.

#### LC-MS for lipidomic analysis

For analysis, the dried lipid extracts were re-suspended in 150 μL mobile phase B (see below) and centrifuged again at maximum speed at 4°C for 5 min. Lipid extracts were analyzed using a Waters ACQUITY I class UPLC system coupled to a mass spectrometer (Thermo Exactive Plus Orbitrap) which was operated in switching positive and negative ionization mode. The analysis was performed using Acquity UPLC System combined with chromatographic conditions as described in (Malitsky et al., 2016) with small alterations. Briefly, the chromatographic separation was performed on an ACQUITY UPLC BEH C8 column (2.1×100 mm, i.d., 1.7 μm) (Waters Corp., MA, USA). The mobile phase A consisted of 45% water (UPLC grade) with 1% 1 M NH4Ac, 0.1% acetic acid, and of 55% mobile phase B (acetonitrile: isopropanol (7:3) with 1% 1 M NH4Ac, 0.1% acetic acid). The column was maintained at 40°C and flow rate of mobile phase was 0.4 mL/min. Mobile phase A was run for 1 min at 100%, then it was gradually reduced to 25% at 12 min, following a gradual decrease to 0% at 16 min. Then, mobile phase B was run at 100% until 21 min, and mobile phase A was set to 100% at 21.5 min. Finally, the column was equilibrated at 100% A until 25 min.

#### Lipid identification and quantification

Orbitrap data was analyzed using LipidSearch™ software (Thermo Fisher Scientific). The validation of the putative identification of lipids was performed by comparing to home-made library which contains lipids produced by various organisms and on the correlation between retention time (RT), carbon chain length, and degree of unsaturation. Relative levels of lipids were normalized to the internal standards and the amount of tissue used for analysis.

#### LC-MS for polar metabolite analysis

For metabolic profiling of the polar phase samples, the lyophilized pellets were dissolved using 150 µL DDW-methanol (1:1), centrifuged twice (at maximum speed) to remove possible precipitants, and were injected into the LC-MS system. Metabolic profiling of the polar phase was performed as described in (Zheng et al., 2015) with minor modifications described below. Briefly, analysis was performed using Acquity I class UPLC System combined with mass spectrometer (Thermo Exactive Plus Orbitrap) which was operated in a negative ionization mode. The LC separation was performed using the SeQuant Zic-pHilic (150 mm × 2.1 mm) with the SeQuant guard column (20 mm × 2.1 mm) (Merck). The Mobile phase B was acetonitrile; Mobile phase A consisted of 20 mM ammonium carbonate with 0.1% ammonia hydroxide in water:acetonitrile (80:20, v/v). The flow rate was kept at 200 μL/min and gradient as follows: 0-2 min 75% of B, 17 min 12.5% of B, 17.1 min 25% of B, 19 min 25% of B, 19.1 min 75% of B, 23 min 75% of B.

#### Polar metabolites identification and quantification

The data processing was performed using TraceFinder™ 4 software (Thermo Fisher) and compounds were identified by accurate mass, retention time, isotope pattern, fragments, and verified using in-house mass spectra library.

#### Mass spectrometry analysis

Samples were normalized by internal standard and sample weight. Metabolites and lipids were omitted if they were detected in less than 70% of the samples (19/27 samples), with a final list of 140 metabolites and 787 lipids. Values were log2 transformed and normalized by the average of each metabolite or lipid. Hierarchical clustering was based on Pearson correlation.

Metabolomics: to perform a statistical analysis we clustered our samples into 4 groups that reflect our experimental design, and performed one-way ANOVA, Tukey post-hoc, and FDR correction. Specifically, experimental groups were clustered according to following (see Figure 3c, left): fully fed (**i**); young fasted (**ii**); old WT fasted (**iii**); and old *APRT^Δ8/+^* fasted (**iv**). We were able to cluster all fully-fed conditions together, as a one-way ANOVA test demonstrated their expression levels were highly similar (i.e. only 4/137 metabolites with a significant difference). Using these clusters, we can estimate a normal response to fasting (comparing clusters i and ii, Tukey post-hoc), and age-dependent deregulated nutrient sensing that is rescued in old *APRT^Δ8/+^* fish (comparing clusters iii and iv, Tukey post-hoc). Significance was called at FDR ≤ 5% for both ANOVA and Tukey post-hoc. These significant metabolites, and the metabolites the hierarchically clustered with them (displaying similar expression patterns) were selected (Figure S3a, blue highlight), and then further clustered according the metabolic pathways BioCyc (Caspi et al., 2020).

Lipidomic: Significant lipids could not be detected, as the lipidomic data was more variable across samples. Additionally, we were not able to treat all fully-fed conditions as a single cluster, as 325/787 lipids (41.2%) displayed significant differences between fully-fed conditions (using ANOVA with FDR correction). Therefore, our statistical test was focused on fasting conditions. We applied one-way ANOVA between 3 clusters: young (**i**); old WT (**ii**); and old *APRT^Δ8/+^* (**iii**). Using these clusters, we estimated age-dependent deregulated nutrient sensing (comparing clusters i and ii) and rejuvenation in old *APRT^Δ8/+^* fish (comparing clusters ii and iii). Significance was called at FDR < 5% for ANOVA, and top 5% for Tukey post-hoc (between groups ii and iii).

### LC-MS for direct quantification of AMP, ADP, and ATP

Metabolites were extracted using standard Methanol/acetonitrile/water, and extracts were snap-frozen. For LC-MS, the Thermo Vanquish Flex ultra-high-performance liquid chromatography (UPLC) system coupled to Orbitrap Exploris 240 Mass Spectrometer (Thermo Fisher Scientific) was used. Resolution was set to 120,000 at 200 mass/charge ratio (m/z) with electrospray ionization and polarity switching mode to enable both positive and negative ions across a mass range of 67–1000 m/z. The chromatography was performed as described previously (Mackay et al., 2015). Briefly, UPLC setup consisted ZIC-pHILIC column (SeQuant; 150 mm × 2.1 mm, 5 μm; Merck). 5 µl of cell extracts were injected and the compounds were separated using a mobile phase gradient of 15 min, starting at 20% aqueous (20 mM ammonium carbonate adjusted to pH 9.2 with 0.1% of 25% ammonium hydroxide): 80% organic (acetonitrile), and terminated with 20% acetonitrile. Flow rate and column temperature were maintained at 0.2 ml/min and 45°C, respectively, for a total run time of 27 min. All metabolites were detected using mass accuracy below 1 ppm. Thermo Xcalibur 4.4 was used for data acquisition. TraceFinder 4.1 was used for data analysis. Peak areas of metabolites were determined by using the exact mass of the single charged ions. The peak areas of different metabolites were determined using Thermo TraceFinder^TM^ 4.1 software, where metabolites were identified by the exact mass of the single charged ion and by known retention time, using an in-house MS library built by running commercial standards of all detected metabolites. For data normalization, raw data files were processed with Compound Discoverer 3.1 to obtain total measurable ions peak intensities for each sample. Each identified metabolite intensity was normalized to the total intensity of the sample. Metabolite-Auto Plotter (Pietzke and Vazquez, 2020) was used for data visualization during data processing.

### RNA sequencing

#### RNA-seq library preparation

Organs were isolated as described above. Samples were disrupted by bead beating in 300 μl of TriZol (Sigma) and a single 3 mm metal bead (Eldan, BL6693003000) using TissueLyzer LT (QIAGEN, #85600) with a dedicated adaptor (QIAGEN, #69980). Beating was performed twice at 50 Hz for 2 min. RNA extraction was performed with Direct-zol RNA Purification Kits (Zymo). RNA concentration and quality were determined by using an Agilent 2100 bioanalyser (Agilent Technologies). Library preparation was performed using KAPA mRNA HyperPrep Kit (ROCHE-08105936001) according to the recommended protocols. Library concentrations were measured by Qubit (dsDNA HS, Q32854), and quality was measured by Tape Station (HS, 5067-5584). Libraries were sequenced by NextSeq 500 high output kit V2, 75 cycle single-end (Illumina, 20024906) using a NextSeq 500 machine (Illumina) with ∼30 million reads per sample.

#### RNA sequencing analysis

Quality control and adapter trimming of the fastq sequence files were performed with FastQC (v0.11.8) (Andrews, 2017), fastx-toolkits (v0.0.13), Trim Galore! (v0.6.4) (Krueger, 2015), and Cutadapt (3.4) (Martin, 2011). Options were set to remove Illumina TruSeq adapters and end sequences to retain high-quality bases with *phred* score > 20 and a remaining length > 20 bp. Successful processing was verified by re-running FastQC. Reads was mapped and quantified to the killifish genome Nfu_20140520 (Reichwald et al., 2015; Valenzano et al., 2015) using STAR 2.7.6a (Dobin et al., 2013). Differential gene expression as a function of age, genotype, and the interaction between age and genotype was performed using the edgeR package (v3.32.1) (McCarthy et al., 2012; Robinson et al., 2010). The three main experimental conditions (males fully fed, males fasted, and female fasted) were sequenced separately. Therefore, analysis was performed on each dataset independently, for increased robustness. Lowly expressed genes were filtered using default parameters in edgeR, and samples were normalized by TMM (Robinson and Oshlack, 2010).

#### Gene Ontology Enrichment Analysis

Enriched Gene Ontology (GO) terms associated with transcripts levels (from either old versus young, heterozygous versus wild type, or Genotype-Age interaction analysis) were identified using Gene Set Enrichment Analysis (GSEA) implemented in R package clusterProfiler (v3.18.1) (Yu et al., 2012). All the transcripts were ranked and sorted in descending order based on multiplication of log2 transformed fold change and –log10(FDR). Note that due to random seeding effect in GSEA, the exact p-value and rank of the enriched terms may differ for each run. These random seeds did not qualitatively affect the enrichment analyses. GO terms were based on human GO annotations from org.Hs.eg.db (v3.13.0) (Carlson et al., 2019) and AnnotationDbi (v1.54.1) (Pagès et al., 2021). Heatmap visualization was perform using ComplexHeatmap (v2.8.0) (Gu et al., 2016) with hierarchical clustering using Pearson correlation. The heatmaps were visualized either by standardized log2 transformed normalized count per million (CPM) with all the associated genes in the GO term (the samples itself or average of each condition) or log2 fold-change between heterozygote versus wild-type or old versus young on GO term genes with fold-change > 1.5 in at least one of the conditions. Venn diagram was plotted by VennDiagram (v1.6.20) R package. Gene Set Variance Analysis (GSVA) was performed using GSVA R package (v1.40.1) (Hänzelmann et al., 2013). Gene sets were selected according to significant GO or KEGG terms, and expanded to related pathways. The gene-by-sample matrix was converted to gene-set-by-sample matrix, and the GSVA score was calculated for gene sets with a minimum of 5 detected genes. Data visualization was performed by the ComplexHeatmap package.

#### Principal component analysis (PCA)

Standardized log2 transformed normalized count per million (CPM) were used as input for principal component analysis (PCA). PCA was performed using autoplot function implemented in R package ggfortify (v0.4.12) and plotted using ggplot2 (v3.3.5).

### Survival, maturity, and growth assays

#### Lifespan measurements

For reproducible lifespan experiments, constant housing parameters are very important (Astre et al., 2022; Dodzian et al., 2018). Following hatching, fish were raised with a following density control: 10 fish in a 1-liter tank for week 1, 5 fish in a 3-liter tank for weeks 2-4. From this point onwards, adult fish were genotyped and single housed in a 1-liter tanks for their remaining lifespan. Plastic plants were added for enrichment. Both male and female fish were used for lifespan experiments, and were treated identically. Fish mortality was documented daily starting week 4. Lifespan analyses were performed using GraphPad Prism for all survival curves with a Kaplan-Meier estimator. To compare the survival curves between different experimental groups, we performed log rank test to examine if the survival curves are significantly different.

#### Maturity and growth measurements

As fish had to be independently evaluated prior to genotyping, housing was slightly different from lifespan experiments, and fish were individually housed in a 1-liter tank starting week 2. For sexual maturity assay (in males only) coloration status of the fish was visually scored according to the onset of tail coloration. For measuring growth, fish were imaged at the indicated timepoints with a Canon Digital camera EOS 250D, prime lens Canon EF 40 mm f/2.8 STM that documented body length. To limit vertical movement during imaging, fish were positioned in a water tank with 3 cm water depth, and images were taken from the top using fixed lighting and height. A ruler was included in each image for accurate scale. Body length was then calculated (using Matlab R2021a), by converting pixel number to centimeters using the included reference ruler.

#### Fertility Analysis

Fish fertility was evaluated according to (Harel et al., 2015). Briefly, 5 independent pairs of fish of the indicated genotypes were placed in the same tank, each consisting of one male and one age-matched female. All breeding pairs were allowed to continuously breed on sand trays, and embryos were collected and counted on a weekly basis for 4 weeks. Unfertilized eggs are easily identified, as they die shortly after egg-laying and the yolk becomes opaque. Results were expressed as a ratio of fertilized eggs per week of egg-lay. The 4-week average of eggs collected for each mating pair was considered as one data point. Significance was calculated using unpaired parametric t test in Prism (GraphPad).

### Histology

Tissues samples were processed according to (Gruenbaum-Cohen et al., 2012; Harel and Tzahor, 2012; Harel et al., 2009, 2012; Nathan et al., 2008; Theis et al., 2010). For paraffine sections, the body cavity of the fish was opened, and following a 72 h fixation in 4% PFA solution at 4°C, samples were embedded in either paraffin or 2-hydroxyethyl methacrylate (Electron microscopy Sciences). Sections of 3-6 μm were stained with hematoxylin and eosin and examined by microscopy. For cryosections, isolated livers were incubated overnight in 4% PFA at 4°C, and then placed in 30% sucrose in DPBS at 4°C for cryoprotection until the livers sank (approximately 12 h). Livers were then embedded in OCT freezing media and placed in −20°C for 12 h, and section of 14 μm were used for downstream applications.

#### Pathological findings

Kidney pathologies were examined according to (Humphrey, 2007; Motamedi et al., 2019). Initial pathology was assessed by presence of tubular lesions (nephrosis), were identified as small vacuoles that appeared in the tubular epithelium. Pathology progression was detected by cell swelling and extensive vacuolar degeneration of the tubular epithelium, in which the interstitium (lymphoid tissue) is infiltrated with lymphocytes. Liver pathologies were identified according to (Di Cicco et al., 2011; Feist et al., 2015; Humphrey, 2007; Sternberg, 1992). Initial pathology was assessed by liver fatty changes, including small droplets in hepatocytes. Pathology progression was detected as steatitis, in which hepatocytes are variable in size and some nuclei vary in size. Ovary pathologies were examined according to (Di Cicco et al., 2011; Feist et al., 2015; Motamedi et al., 2019). Briefly, atresia was considered as an increase in degradation and resorption of oocytes in development. Egg debris were detected by the presence of yolk within the oviduct (1,3,4).

#### Staining of neutral lipid droplets

Cryosections were stained for lipids droplets according to (Griffett et al., 2013). Briefly, slides were incubated in 2 μg/mL BODIPY 493/503 (Invitrogen™) in 1X DPBS for 15 min. Sections were washed three times in cold 1X DPBS and then counterstained with 1 μg/mL Hoechst. Three sections per liver, from a total of 3-4 fish, were imaged using the FV-1200 confocal microscope (Olympus, Japan). BODIPY intensity was normalized by nuclear staining (Parafati et al., 2018), quantified by ImageJ, and values were analyzed with GraphPad Prism.

### Dietary manipulations

#### 24 h fasting

Following the morning feeding, fish were fasted for 24 hours.

#### Lifelong intermitted fasting

Starting on week 4, fish were fed once a day for their remaining lifespan with a combination of GEMMA Micro 300 Fish Diet (Skretting Zebrafish, USA) supplemented by live Artemia.

#### Short-term high-fat diet

Old fish (15 weeks old) were fed twice a day, for a total of one week, with BioMar fish diet (0.5mm INICIO Plus SEA BREAM, BioMar Group, Demark), supplemented by live Artemia. This protocol was adapted from a recent report that demonstrated that killifish fed with BioMar INICIO had more visceral fat, and their livers possessed more lipid droplets (Žák et al., 2020). As demonstrated for other short-term approaches in zebrafish (Landgraf et al., 2017), one week of feeding with BioMar was sufficient for inducing significant accumulation of lipid droplets in the liver (when compared with our normal feeding, Figure S7g).

#### Antibodies validation

To validate antibody specificity using western blot, we generated two conditions in which AMPK-related pathways are expected to be altered. Specifically, male killifish were either starved for 3 days (Mohapatra et al., 2020) or exposed to 20 mM 2-deoxy glucose for 2.5 h in system water (2-DG, Sigma Aldrich) (LANE et al., 1998).

#### 17β-estradiol (E2) exposure

17β-estradiol (E2, Holland Moran) was dissolved in ethanol, and sprayed on food pellets at 100 mg/kg for both drugs (Mohapatra et al., 2020). Food was then dried overnight under a chemical fume hood. Adult fish (9 weeks old) were individualized in tanks containing system water, and fed twice a day for a period of 8 days. Control groups were fed with ethanol treated food pellets and were similarly maintained and sampled. As expected, in response to the 17β-estradiol treatment, we observed lipid accumulation in treated fish (Cakmak et al., 2006).

### Generation of primary fibroblast cultures from killifish tail fins

Adult fish (8-12 weeks old), of the indicated genotype and gender were sedated with MS-222 (200 mg/L Tricaine, in system water). All following experiments were conducted at 28°C unless stated otherwise. Following sedation, a 2–3 mm tissue was trimmed from the tail fin using a sterile razor blade, and individually disinfected for 10 min with a 25 ppm iodine solution (PVP-I, Holland Moran 229471000) in DPBS (Biological Industries). Followed by a rinse with DPBS, tissue samples were incubated for 2 h with 1 mL of an antibiotic solution containing Gentamicin (50 µg/mL Gibco) and Primocin^TM^ (50 µg/mL, InvivoGen) in DPBS at room temperature. Tissues were then transferred into an enzymatic digestion buffer (200 µL, in a 24-well plate) containing Dispase II (2 mg/mL, Sigma Aldrich) and Collagenase Type P (0.4 mg/mL, Merck Millipore) in Leibovitz’s L-15 Medium (Gibco). During the first 7 days, cells were daily washed with fresh media before adding new media. When cells reached 85– 90% confluency, they were passaged with Trypsin-EDTA 0.05% (0.25% Trypsin-EDTA, diluted in DPBS). Cells were incubated at 28°C humidified incubator (Binder, Thermo Scientific) with normal air composition, and were used for downstream applications between passages 4-12.

### Seahorse Metabolic Extracellular Flux Profiling

The Seahorse XFp Analyzer (Agilent Technologies, USA) was used to measure oxygen consumption rates (OCR, in pmol O_2_ per min), and extracellular acidification rates (ECAR, in mpH per min). Primary fibroblasts from killifish tail fins were cultured as described above. The day before the experiment, cells were seeded (50,000 cells/well) in XFp 8-well plates (Agilent Technologies, USA). A relatively large number of cells was used in our assay to compensate for the small size and low basal respiration of primary fish cells. All the required drugs were supplied by the manufacturer (Seahorse Bioscience), and drug injection times were according to the XFp standard protocol. OCR and ECAR measurements were analyzed according to manufacturer’s instructions and GraphPad Prism (v7, GraphPad).

Although cells were seeded in equal numbers to facilitate OCR and ECAR normalization, actual number of cells in each well was further confirmed using imaging to account for potential differences in proliferation rates. Briefly, following each metabolic assay, cells were fixed and stained for 15 min with 4% paraformaldehyde and 10 µM Hoechst 33342 (Thermo Fisher) in DPBS. Each well was imaged three times (IX83, Olympus), and averaged cell density (number of cells per field of view) was counted using CellProfiler (https://cellprofiler.org). Finally, the total number of cells per well was estimated by the relative area represented by the field of view, compared to the total area of the well. The following assays were performed 2-3 times. Each experiment contains at least three independent samples per genotype.

#### Cell Mito Stress Assay

Cell Mito Stress Test (XFp Cell Mito Stress Test Kit, Agilent Technologies) was performed following the manufacturers guidelines, with minor adaptations for fish cells described below. 1 h before the measurements, culture medium was replaced and the cells were incubated for 1 h min at 28°C with the Seahorse XF Base medium (Seahorse Bioscience), supplemented with 2 mM L-glutamine and 1 mM pyruvate (Seahorse Bioscience). Instead of glucose, we used 5 mM galactose (Sigma-Aldrich), which is the sugar source used by the L15 culture media. Oxygen consumption rate (OCR) and Extracellular Acidification Rate (ECAR) were detected after injection of oligomycin (1 μM), Carbonyl cyanide-p-trifluoromethoxyphenylhydrazone (FCCP, 1 μM), and the combination of rotenone & antimycin A (Rot/AA, 1 μM).

#### Cell Glycolysis Stress Assay

Cell Glycolysis Stress Test (XFp Glycolysis Stress Test Kit, Agilent Technologies) was performed following the standard protocol. Culture media was replaced by Seahorse XF Base medium, supplemented with 2 mM L-glutamine. During the test, final concentration of 15 mM glucose, 1 μM oligomycin, and 50 mM of 2-deoxyglucose (Seahorse Bioscience) were injected according to the XFp standard protocol.

#### Proliferation of cultured cells

Cells from individual fish were seeded in 60 mm dish in triplicates, at a density of 200,000 cells/well. 24 h, 48 h, 72 h after seeding cells were counted using Neubauer counting chamber (MarienField #0640110). Medium was changed 4 h before the first counting point. Experiment was performed twice using 3 independent replicates per genotype (using the mean of three measurements per dish). Significance was calculated using two-way ANOVA test and corrected by two-stage linear step-up procedure of Benjamini, Krieger and Yekutieli.

### Pharmacological manipulation of cultured cells

#### Adenine treatment

Cells were seeded in XFp 8-well plates (Agilent Technologies, USA) at a density of 50,000 cells/well and cultured overnight with adenine 10 µM. 1 h prior to the seahorse experiment, culture medium was replaced with the Seahorse XF Base medium (Agilent Technologies, USA) supplemented with adenine 10 µM (Sigma Aldrich), and cells were incubated for 1 h min at 28°C.

#### 8-azaadenine treatment

To characterize and validate the relative loss of APRT activity in heterozygous cells, an adenine analog, 8-azaadenine (Holland Moran) was used according to (Jones and Sargent, 1974). Briefly, mutant APRT cells that have reduced ability to metabolize adenine exhibit partial resistance to the toxic intermediate generated by AA (Jones and Sargent, 1974). Cells were treated with a range of concentrations of AA (0-200 µg/mL) for 4 days without media change. As AA is dissolved in 1 M NaOH, an equal amount of 1 M NaOH was added to the controls. Experiments were performed twice using 3 independent replicates per genotype.

#### Serum starvation

Cells were seeded in a 12-well plate at a density of 100,000 cells/well. The following day, the medium was replaced with fresh media for 4 h, with L-15 medium containing either or 15% (control) or 5% FBS.

### Mitochondrial morphology

#### Fluorescence microscopy

Cells were plated on a µ-Slide 8 Well Glass Bottom plates (ibidi, #80827), at a density of 40000 cells/well. Confocal imaging was performed on live cells using a sealed, environmentally controlled chamber without CO_2_ at 28°C. A Day after, cells were labeled with 200 nM Mitoview 633 (Biotium) for 15 min prior to live imaging. Labeled cells were imaged on a laser scanning confocal microscope (FV-1200, Olympus, Japan) using a 60X/1.42 oil immersion objective. Scans were acquired using a sequential mode. All fluorophores were excited on separate tracks. Hoechst was excited with 405 nm and emission was captured through 485 nm short-pass filter. Mitoview 633 was excited with 561 nm and emission was captured through a 570-620 nm.

#### Mitochondrial footprint and network analysis

To analyze mitochondrial morphology, we used MiNA (http://github.com/ScienceToolkit/MiNA), a plug-in macro toolset for Fiji (Schindelin et al., 2012). This workflow estimates mitochondrial footprint from a binarized copy of the provided image, and the lengths of mitochondrial networks are estimated using a topological skeleton (mitochondrial length was defined as the length of rods and network branches). Additionally, we used the generated overlays (or a 3D rendering) to further confirm the accuracy of the analysis. Experiment were performed twice using 3 independent replicates per genotype. Three field of views, containing between 4-8 cells, were independently analyzed as described above.

### Protein Gel Electrophoresis and Immunoblotting

#### Liver samples

Individual killifish, according to the specified age, gender, genotype, and feeding condition, were euthanized in 400 mg/L of Tricaine in system water. Animals were dissected on ice under a stereo microscope (Leica S9E) according to (Astre et al., 2022). All procedures were carried at 4°C unless stated otherwise. Homogenization was performed in a 2 mL Safe-Lock Tubes (Eppendorf), to allow full homogenization of liver tissue, using 3 mm-metal beads (Eldan Israel, Cat# RC55 420) and RIPA lysis buffer (7.9 g/L Tris-HCl, 9 g/L NaCl, 0.76 g/L EGTA, 10 ml 10% Triton X-100, pH 7.2). Prior to homogenization, 200 µl buffer was added to ∼half a liver, and freshly supplemented with anti-protease and anti-phosphatase cocktail (Biotool). Homogenization was carried out with mechanical disruption using TissueLyzer LT (QIAGEN, #85600) with a dedicated adaptor (QIAGEN, #69980) at 50 Hz for 2 min X2. Protein concentration was measured with Pierce™ BCA Protein Assay Kit (Thermo Scientific™), according to manufacture instructions. 5-10 µg of liver homogenate was then mixed with the Sample Buffer (Tris-HCl pH 6.8 62.5 mM, Glycerol 10%, SDS 2%) at a ratio of 1:3. 2-mercaptoethanol (Sigma Aldrich) (5%) and Bromophenol blue (Sigma Aldrich) (0.02%) were freshly added, boiled at 95°C for 10 min, and placed on ice.

#### Primary culture

Following a wash with cold DPBS, cells were re-suspended with 250 µl of the Sample Buffer (Tris-HCl pH 6.8 62.5 mM, Glycerol 10%, SDS 2%) at a ratio of 1:3, freshly supplemented with 5% 2-mercaptoethanol (Sigma Aldrich), 0.02% Bromophenol blue (Sigma Aldrich), and anti-protease and anti-phosphatase cocktail (Biotool). Samples were then boiled at 95°C for 10 min. 10 µl were used for western blot.

#### Immunoblotting

Standard western blot was performed. Briefly, protein extracted (5-10 µg from livers, or 10 µl from cells) were resolved using Novex WedgeWell 4-20% Tris-Glycine gel (XP04205BOX, Thermo Fisher) and electrophoresed in Cell SureLock (Novex) in constant voltage of 50 V for 15 min (to clear stacking), followed by 120 V for 1 h. Transfer to a Nitrocellulose membrane was performed using the iBlot 2 (Thermo Scientific, IB21001), followed by Ponceau Red red staining (Sigma Aldrich, P7170-1L). Membranes were washed in TBST 0.1% (Tris-HCl 10mM pH 8.0, NaCl 150 mM, Tween 0.1%) once for 5 min to remove Ponceau staining, and blocked in 5% BSA (Sigma-Aldrich) in TBST for 1 h at room temperature. The resolved proteins were probed with the following antibodies at a concentration of 1:1000 (diluted in blocking solution at 4°C overnight): anti-AMPKα (CST-5831), anti-Phospho-AMPKα (Thr172) (CST-2535), anti-ACC (CST-3676), anti-Phospho-ACC (CST-3661), anti-S6 Ribosomal Protein (CST-2217), anti-Phospho-S6 Ribosomal Protein (CST-2211), and anti-Actin (MP Bio 0869100). Following three washes of 10min, membranes were incubated with 1:5000 HRP-conjugated goat anti-rabbit (ab6721, Abcam) or goat anti-mouse (ab6789, Abcam) antibodies for 1 h, and washed three times for 10 min. Chemiluminescence was detected using EZ-ECL kit (Biological Industry) and imaged with either ChemiDoc MP Imaging System (Biorad) or Fusion Pulse 6 (Vilbert Loumat). Band densitometry was quantified using ImageJ, and normalized according to Actin values.

### DNA isolation and measurement of mitochondrial DNA content using quantitative PCR

Quantification was performed according to (Hartmann et al., 2011). Briefly, total DNA from cells, tails and liver was extracted using QIAamp Micro kit (Qiagen) according to the manufacturers protocol. The relative mitochondrial DNA (mtDNA) copy number was determined by quantitative PCR (qPCR) with the nuclear *CDKN2A/B* gene and the mitochondrial *16S rRNA* gene (primer sequences are available at (Hartmann et al., 2011)). qPCR was performed with Magnetic Induction Cycler (Mic, bio molecular systems) using Fast SYBR Green Master Mix (2X) (ThermoFisher 4385610), 100 nM of each primer, and 20 ng DNA as template. All reactions included negative controls (without template, or without the enzyme). Ct values of mtDNA were normalized to Ct values of the nuclear locus according to (Hartmann et al., 2011), using the following equation: relative mtDNA copy number per diploid cell= 2×2^ΔCt^, where ΔCt is Ct_CDKN2A/B locus_−Ct*_16S rRNA_* _locus_.

### A-to-I RNA editing

A-to-I editing is the most prevalent RNA editing. It is performed by double-stranded RNA-specific adenosine deaminase (ADAR) family of proteins (Eisenberg and Levanon, 2018). Sequencing machines recognize Inosine as Guanosine, thus allowing us to quantify A-to-I editing by counting mismatches from Adenosine to Guanosine (A2G)^2^.

#### Quality control for RNA editing

Additional filtration steps, in addition to those performed for differential gene expression, were taken to eliminate common biases in RNA Editing. Specifically, we used PRINSEQ-lite 0.20.4 (Schmieder and Edwards, 2011) to remove exact duplicates, retain reads no shorter than 72 bp, and trim reads longer than 63 bp. Duplicate RNA reads, defined as reads with the same sequence on the same strand or with the reverse-compliment sequence on the opposite strand, can result from PCR cycles conducted before sequencing. Using only reads with similar lengths allowed us to compare the RNA Editing Index of different samples. Trimmomatic 0.39 (Bolger et al., 2014) was used to remove Nextera transposase adapters. Finally, we assessed the quality of the reads with FastQC 0.11.8 (Andrews, 2017) and MultiQC 1.11 (Ewels et al., 2016).

#### Genome and genomic annotations

Reads was mapped and quantified to the killifish genome Nfu_20140520 (Reichwald et al., 2015; Valenzano et al., 2015). As ADAR target editing sites are usually prevalent in repetitive regions (Eisenberg and Levanon, 2018), we created a comprehensive annotation of these regions in the killifish genome by expanding the initial annotation. Specifically, we extracted the RepeatMaskerLib.embl library database using RepeatMasker 4.1.0 (Chen, 2004), and identified coding regions using the available GFF file. Finally, we used EDTA 1.9.6 (Ou et al., 2019) to create an extended library of repeats.

#### Signal-to-noise ratio (SNR)

Considering A-to-G editing as *signal* and the 2^nd^ most prevalent DNA-RNA mismatch other than A-to-G as *Noise*, we define SNR as:

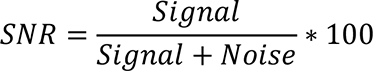

#### Hyper editing & cluster screening

ADAR’s activity is characterized by dense clusters of editing sites, which results in reads that are difficult to align by regular alignment procedures. Therefore, we applied an adaptation to successfully align those hyper-edited reads to the genome (Porath et al., 2014). The output of this method is a list of hyper-edited sites and ultra-edited regions (UE). To get a clear SNR as possible, we applied a previously published approach (Buchumenski et al., 2021), termed here as Cluster Screening. Using a distance of 20 bp, we first required that two different editing sites (e.g., A2G and C2T) cannot reside within 20 bp next to each other. Second, each editing site must have a neighbor editing site of the same type, located no further than 20 bp from him. The first demand is meant to overcome alignment errors that are the result of duplication events. The second demand helps to identify clusters of editing sites, contrary to SNPs that occur randomly at various genomic locations. We dismissed editing sites that fail to satisfy either of these requirements.

#### RNA editing index

To quantify the global editing levels of each UE cluster, we ran the RNA Editing Index tool

(Roth et al., 2019). For each kind of mismatch from *Reference Base* to *Mutated Base*, we define the Index of a region as

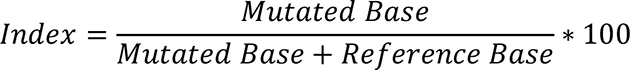

where *Reference Base* is the number of reads mapped to genomic positions (of that base), and *Mutated Base* is the number of reads with *Reference Base*to **Mutated Base** mismatch mapped to those positions. Specifically, the Index is a weighted average of RNA editing levels of a mismatch across a region. Using the Index allows us to compare RNA editing of different regions and samples. We ran the RNA Editing Indexer (https://github.com/a2iEditing/RNAEditingIndexer) with the following parameters as input: (1) The average expression level of each gene, as quantified by Salmon v1.4 (Patro et al., 2017); (2) RefSeq annotations of the genes and aligned BAM files. Independently of the Hyper Editing tool alignment, we aligned the Fastq files to the genome using BWA 0.7.17 (Li and Durbin, 2009) with the MEM algorithm; (3) The UE clusters we previously chose with Cluster Screening. We ran the Index both in stranded and unstranded mode, but as the SNR of the stranded runs was lower (data not shown), we decided to use its unstranded output. Independent t-tests were done using scipy.stats.ttest_ind 1.6.2, multiple independent t-tests were corrected using statsmodels.stats.multitest.fdrcorrection 0.12.2 using default parameters.

---

**Figure S1:**
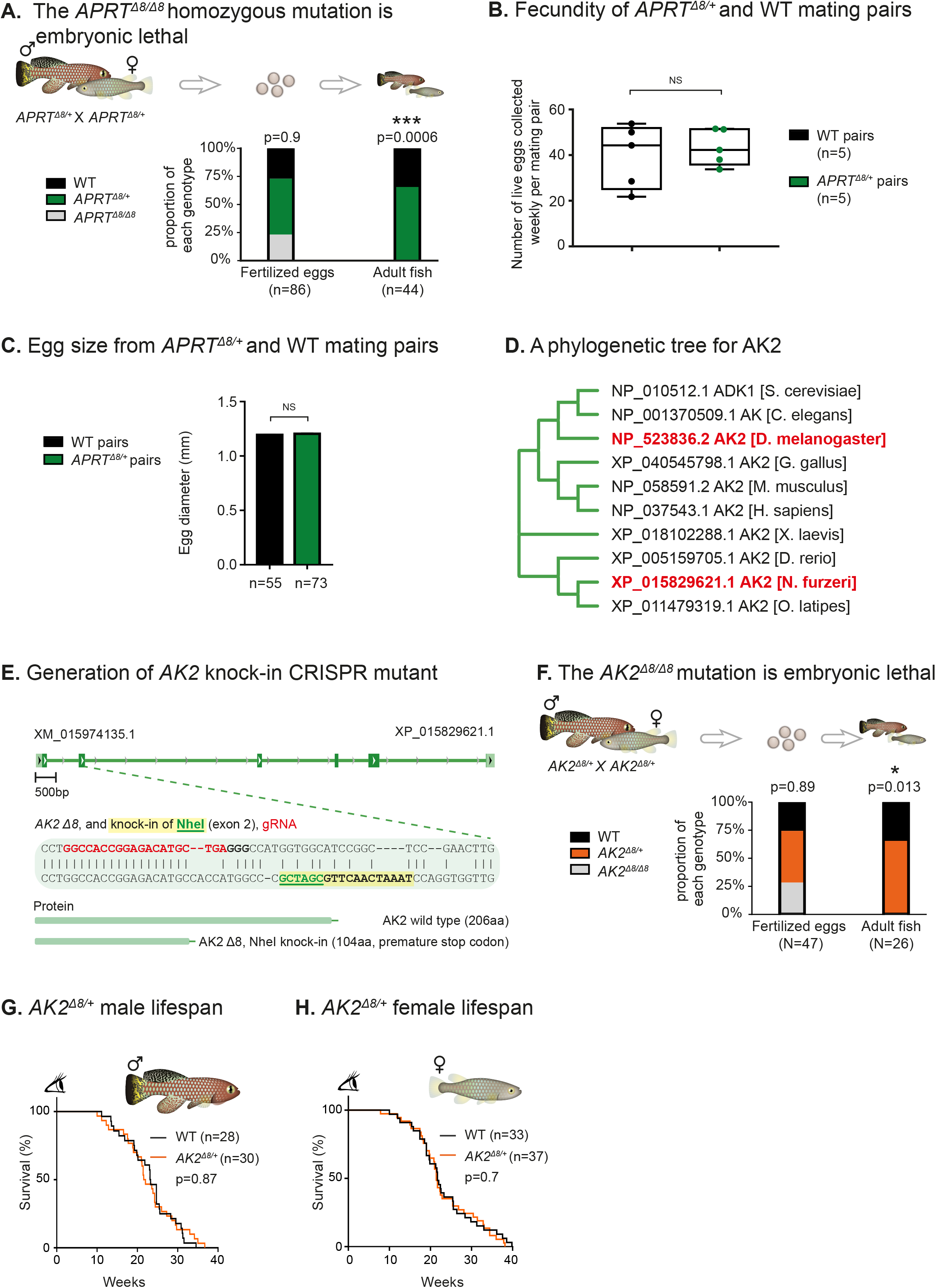
Male and female lifespan is unaffected by the *AK2^Δ8/+^* heterozygous mutation. (**A**) Mating of *APRT^Δ8/+^* heterozygous pairs to generate homozygous *APRT^Δ8/Δ8^* mutants. *APRT^Δ8/Δ8^* fertilized eggs are observed at the expected Mendelian ratios (no difference between expected and observed frequencies). *APRT^Δ8/Δ8^* homozygous adult fish are not detected, indicating embryonic lethality. Significance was calculated using χ^2^ test. (**B**) Number of live eggs for each mating pair of the indicated genotypes. Each data point is an average of 4 weekly embryo collections from a single mating pair (a total of 5 pairs per genotype). Bars represent minimum and maximum. Significance was calculated using an unpaired t-test. (**C**) Egg size from mating pairs of the indicated genotypes (size is measured as egg diameter, in millimeter). Total number of eggs per experimental condition is presented. Bars represent mean ± SEM (due to low variation, bars are very small). Significance was calculated using an unpaired t-test. (**D**) A phylogenetic tree for adenylate kinase 2 (AK2) protein sequence for classical genetic model systems. A neighbor-joining tree with no distance corrections (Clustal Omega, EBI). (**E**) Generation of *AK2* CRISPR mutant, including guide RNA (gRNA) target design for exon 2 (red sequence), knock-in of a NheI restriction site (green), and successful germline transmission of an 8 bp deletion (Δ8). The *AK2* Δ8 allele is predicted to generate a protein with a premature stop codon. (**F**) Mating of *AK2^Δ8/+^* heterozygous pairs to generate homozygous *AK2^Δ8/Δ8^* mutants. *AK2^Δ8/Δ8^* fertilized eggs are observed at the expected Mendelian ratios (no difference between expected and observed frequencies). *AK2^Δ8/Δ8^* homozygous adult fish are not detected, indicating embryonic lethality. Significance was calculated using χ^2^ test. (**G-H**) Lifespan of WT and *AK2^Δ8/+^* fish performed separately for males (G) and females (H) p-values for differential survival in log-rank tests, and fish numbers are indicated. See Table S1 for additional information.

**Figure S2:**
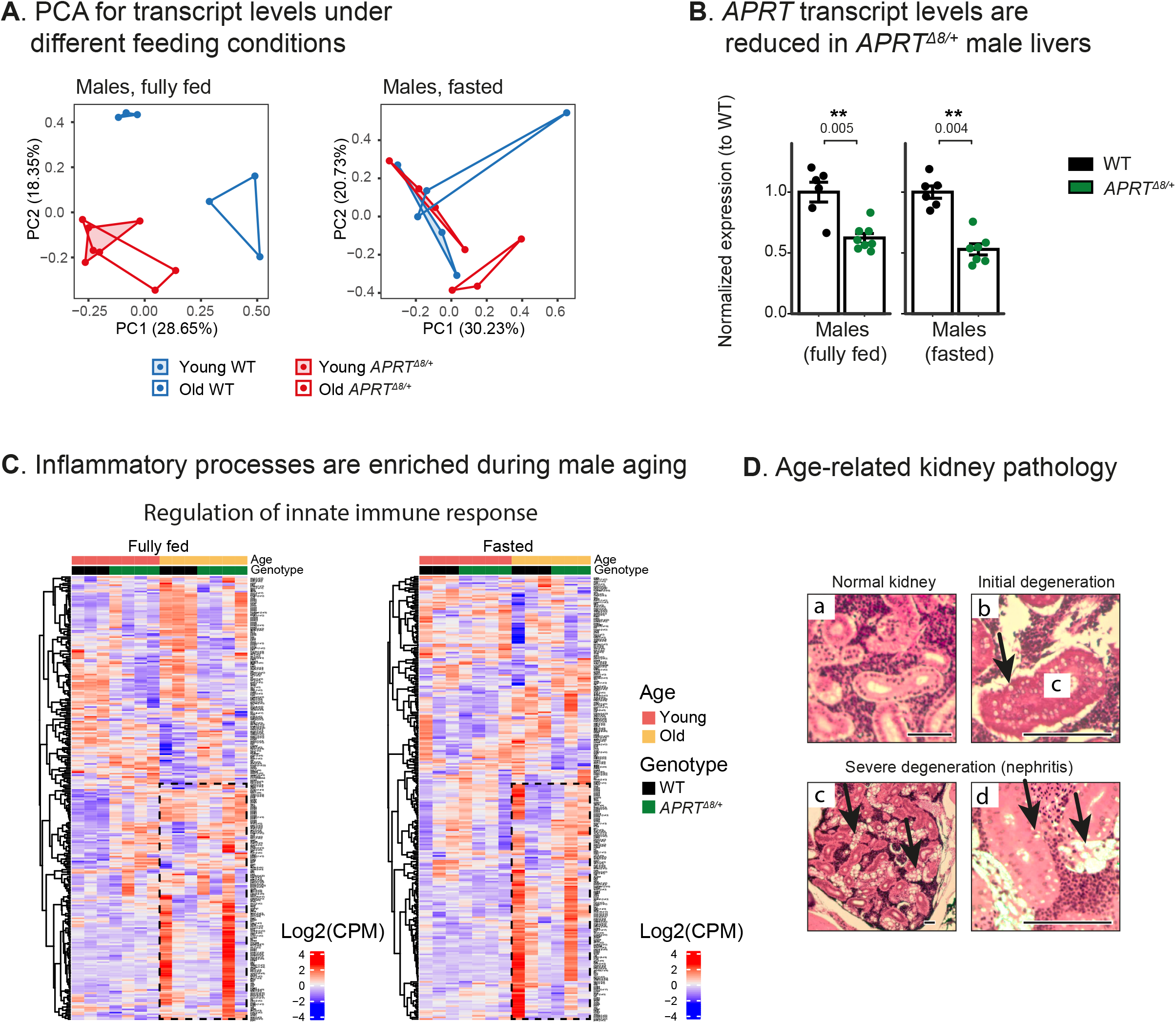
Transcriptomic analysis for male fish. (**A**) Principal component analysis (PCA, for PC1 and PC2) for transcript levels from the livers of WT (blue) or *APRT^Δ8/+^* (red) fish, young (shaded) or old (nonshaded). Experiments were performed under fully-fed (top) or fasting conditions (bottom). Each symbol represents an individual fish. (**B**) Relative expression of *APRT* mRNA in WT and *APRT^Δ8/+^* fish using RNA-seq. For each condition, expression level is normalized to WT. Each symbol represents an individual fish. Experimental groups differ by genotype and feeding condition (old and young fish are grouped together). Significance was calculated using an unpaired t-test, with FDR correction. (**C**) Heatmap for inflammation-related gene expression in male livers (GO term: regulation of innate immune response). Genes are hierarchically clustered, and expression values are presented as z-score on Log2(CPM). Each column represents an individual fish at the indicated experimental conditions. Dashed lines highlight differences between young and old fish. (**D**) Histological characterization of age-related kidney degeneration (black arrows). a) Normal kidney. b) First signs of degeneration, with vacuoles in the tubular epithelium. c-d) Severe degeneration, nephritis, with severe vacuolation in tubules (c) and vacuolar changes of renal tubules (d). Images are from either young (a) or old (b-d) males (n=3 for each group). Scale bar 200 µm.

**Figure S3:**
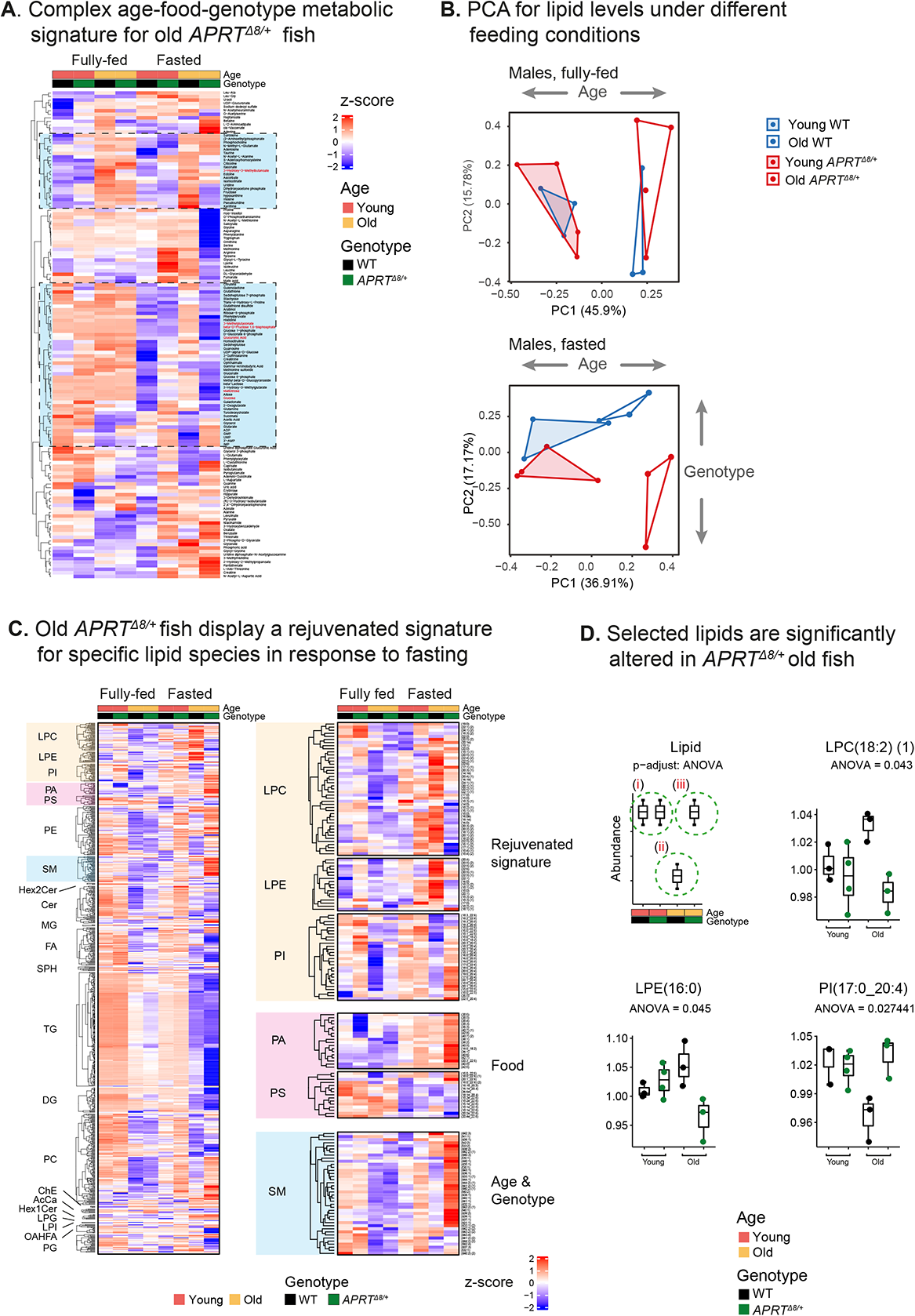
Liver metabolomic and lipidomic analysis by age, genotype, and feeding condition in male fish. (**A**) Hierarchical clustering of all identified liver metabolites. Each square represents normalized relative abundance (indicated by color) from an average of 3-4 fish of the indicated experimental condition. Each condition differs by age (young or old), genotype (WT or *APRT^Δ8/+^*), and feeding condition (fully-fed or fasted for 24 h). Significant metabolites are in red, and a broader list of selected metabolites that cluster with them are highlighted in blue. Statistical tests as described in Figure 3b. Selected metabolites (blue) were further analyzed in Figure 3b, 3c. (**B**) Principal component analysis (PCA) for abundance of lipid species from the livers of WT (blue) or *APRT^Δ8/+^* (red) fish, young (shaded) or old (nonshaded). Experiments were performed under fully-fed (top) or fasting conditions (bottom). Each symbol represents an individual fish. Full metabolite list is reported in Table S3. (**C**) Heatmap showing all identified (left) and selected (right) lipid species from the livers of the indicated experimental groups. Specific glycerophospholipids (i.e. PI, LPE, and LPC) display a restoration of youthful levels in old *APRT^Δ8/+^* fish under fasting conditions (right). Other lipid species display a complex dependency on diet (i.e. PA, PS), or age and genotype (i.e. SM). Each square represents normalized relative abundance (indicated by color), from an average of 3-4 fish. LPC: Lysophosphatidylcholine; LPE: lysophosphatidylethanolamine; PI: Phosphtatidylinositols; PA: Phosphtatidic acids; PS: Phosphtatidylserines; PE: Phosphtatidylethanolamines; SM: Sphingomyelins; Hex1Cer/Hex2Cer: hexa-ceramides 1/2 glycan chains; Cer: Ceramides/Sphingoid bases; MG: Mono(acyl|alkyl) glycerols; FA: Fatty acids; SPH: Ceramides/Sphingoid bases; TG: Tri(acyl|alkyl) glycerols; DG: Di(acyl|alkyl) glycerols; PC: Phosphtatidylcholines; ChE: Cholestrol ester; AcCa: acylcarnitine; LPG: Lysophosphatidylglycerol; LPI: Lysophosphatidylinositol; OAHFA: (O-Acyl)-ω-hydroxy fatty acids; PG: Phosphtatidylglycerols. Full lipid list is reported in Table S3. (**D**) Boxplot showing trending specific lipid species selected from (**C**). Each dot represents one fish. Bars represent minimum and maximum. Significant lipids could not be detected due to sample noise. Therefore, our statistical test was limited to fasting conditions. We applied one-way ANOVA between 3 clusters: young (**i**); old WT (**ii**); and old *APRT^Δ8/+^* (**iii**). Using these clusters, we estimated age-dependent deregulated nutrient sensing (comparing clusters **i** and **ii**) and rejuvenation in old *APRT^Δ8/+^* fish (comparing clusters **ii** and **iii**). Significance was called at FDR < 5% for ANOVA, and top 5% for Tukey post-hoc (between groups **ii** and **iii**).

**Figure S4:**
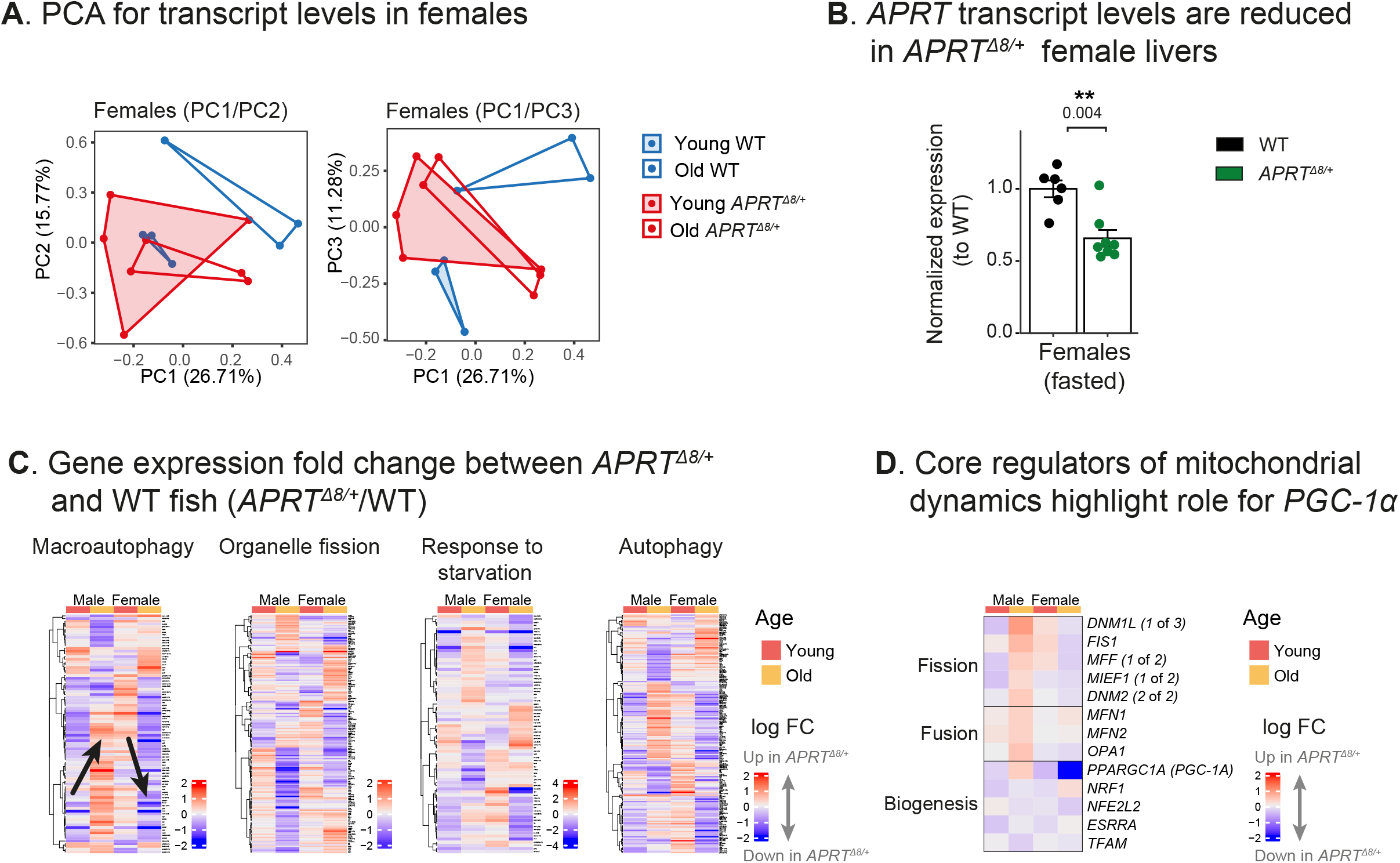
Transcriptomic analysis for female fish. (**A**) Principal component analysis (PCA) for transcript levels from the livers of WT (blue) or *APRT^Δ8/+^* (red) female fish, young (shaded) or old (nonshaded). Experiments were performed under fasted conditions. Each symbol represents an individual fish. PC1/PC2 and PC1/PC3 are shown. (**B**) Relative expression of *APRT* mRNA in WT and *APRT^Δ8/+^* fish using RNA-seq. For each condition, expression level is normalized to WT. Each symbol represents an individual fish. Experimental groups differ by genotype (old and young fish are grouped together). Significance was calculated using an unpaired t-test, with FDR correction. (**C**) Heatmap for genotype-related gene expression changes (log2 fold change) between *APRT^Δ8/+^* and WT expression values using a cutoff of 1.5-fold change. Experimental groups differ by sex (male or female) and age (young or old). Selected pathways are from Figure 4d, and black arrows highlight age-related changes in gene expression. (**D**) Heatmap for genotype-related gene expression changes (log2 fold change) between *APRT^Δ8/+^* and WT expression values depicting core regulators of mitochondrial fusion, fission, and biogenesis. Experimental groups differ by sex (male or female) and age (young or old).

**Figure S5:**
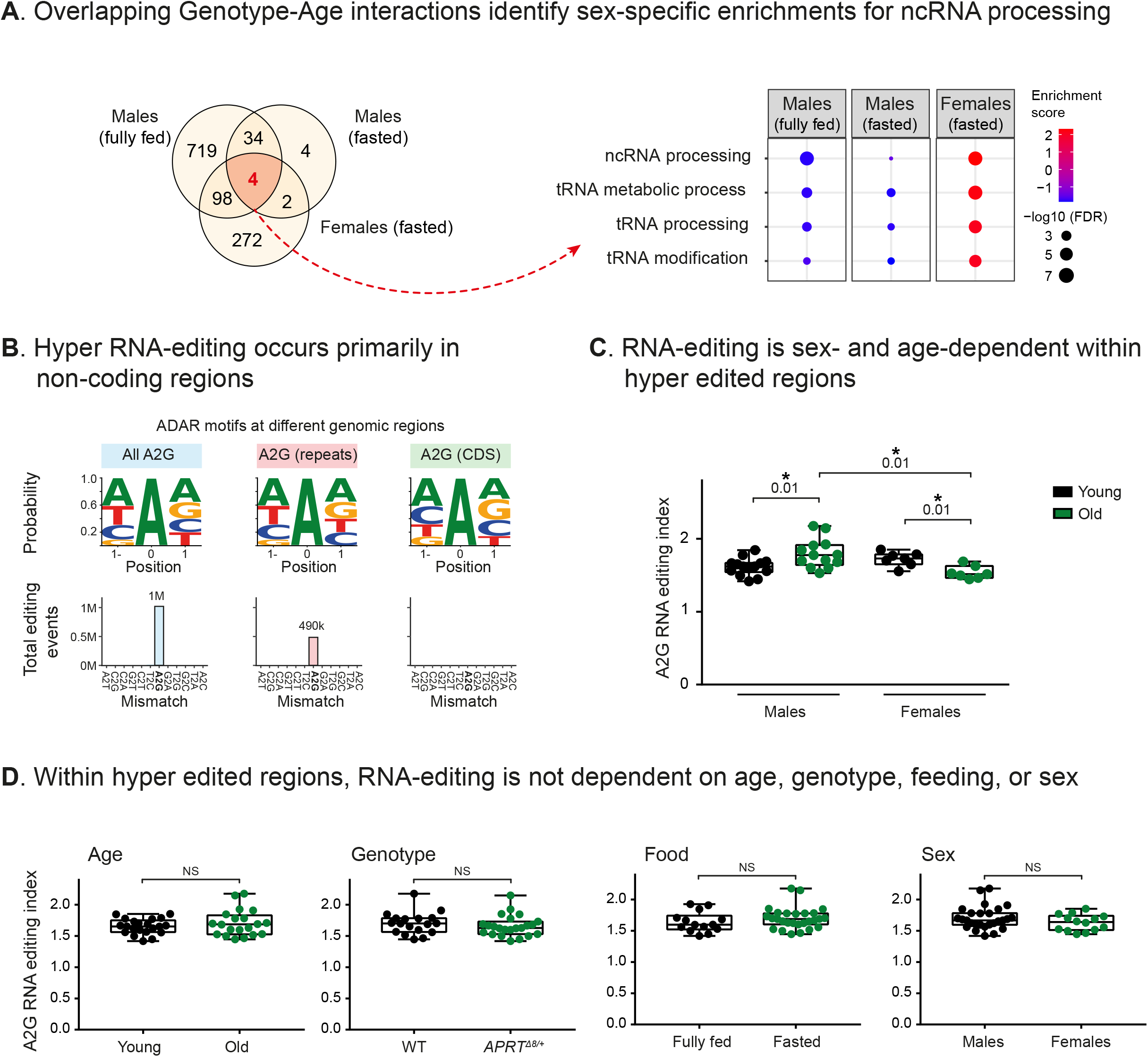
RNA-editing is sex- and age-dependent. (**A**) Venn diagram showing the overlap of significantly enriched pathways that were identified by Genotype-Age interaction analysis across all experimental groups (left). Analysis was performed separately for each experimental condition (i.e. male fully-fed, male fasted, and female fasted). GO enrichments using GSEA were called at FDR < 5% (right). (**B**) Identification of A-to-I hyper editing in killifish at different genomic regions across all experimental groups. Signal-to-noise was assessed by estimating the nucleotide frequencies at the hyper edited sites (that matched the ADAR deamination motif, top), and by the number A- to-G mismatches compared to all types of mismatches (bottom). (**C**) Stratifying RNA-editing by experimental conditions highlights significant sex- and age-dependency within hyper edited regions. Additional comparisons are available in Figure S5d. Each symbol represents an individual fish. Bars represent minimum and maximum, and significance was measured by multiple independent unpaired t-test with FDR correction. (**D**) Stratifying RNA-editing by experimental conditions. Each symbol represents an individual fish. Bars represent minimum and maximum. Significance was measured by an independent unpaired t-test, and exact p-values are indicated in Table S4.

**Figure S6:**
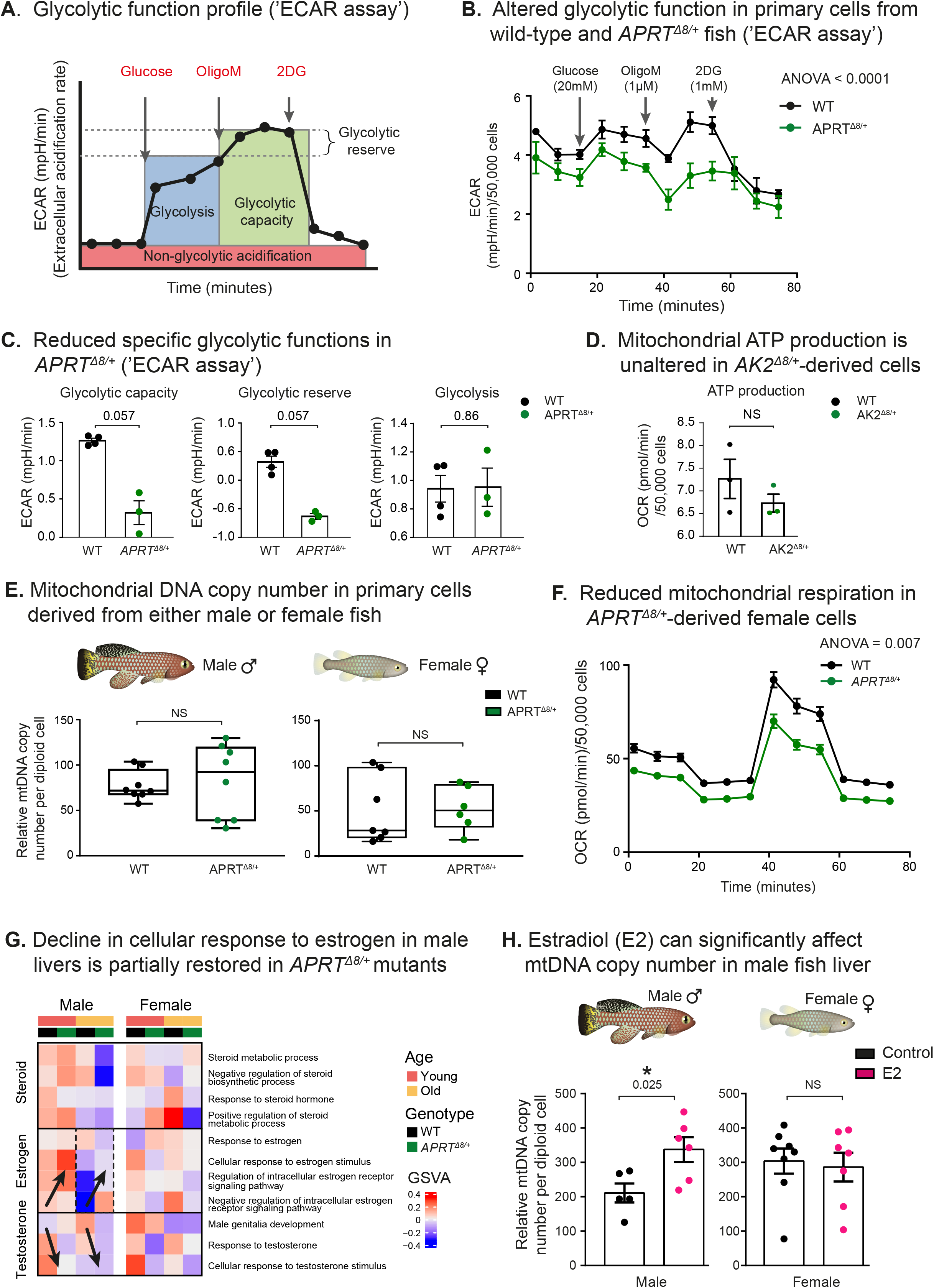
Characterization of mitochondrial functions and copy number. (**A**) Measurement of glycolytic function profile by extracellular acidification rate (ECAR assay, extracellular acidification rate in mpH/min) using the Seahorse Glycolysis Stress Test Kit (Agilent). Measurements were performed under basal conditions, or following the addition of glucose, oligomycin (OligoM, an ATP synthase inhibitor), or 2-deoxy glucose (2-DG, a glucose analog, that inhibits glycolysis). Specific parameters were further normalization by actual cell numbers. n = 3-4 for each genotype. (**B**) Measuring glycolytic function in WT- and *APRT^Δ8/+^*-derived primary cells. Each symbol represents an average of three independent measurements using cells derived from 3-4 individual fish. Error bars represent mean ± SEM, and significance was measured using two-way ANOVA and BKY correction. (**C**) Glycolytic capacity (left), Glycolytic reserve (center), and Glycolysis (right) were compared between WT- and *APRT^Δ8/+^*-derived primary cells. Each symbol represents an individual fish. Error bars represent mean ± SEM, significance was measured using Mann-Whitney test, and exact p-values are indicated. (**D**) ATP production was compared between WT- and *AK2^Δ8/+^*-derived primary cells. Each symbol represents an individual fish. Error bars represent mean ± SEM, and significance was measured using Mann-Whitney test. (**E**) Relative mtDNA copy number was determined by quantitative real-time PCR (qPCR) using primers for mitochondrial (*16S rRNA*) and nuclear (*CDKN2A/B*) gene. Each symbol represents a primary culture isolated from an individual male fish of the indicated genotype. Bars represent minimum and maximum and significance was estimated by unpaired t-test. (**F**) Measuring mitochondrial respiration in WT- and *APRT^Δ8/+^*-derived primary cells, isolated from female fish. Each symbol represents an average of three independent measurements using cells derived from three individual fish. Error bars represent mean ± SEM, and significance was measured by two-way ANOVA and BKY correction. (**G**) Heatmap for gene expression changes of steroid and steroid hormones related pathways (GO), in male and female livers using GSVA, highlighting age- and genotype-dependent signatures. Arrows indicated overall increase (up) or decrease (down) in GSVA score. (**H**) Relative mtDNA copy number in adult fish livers was determined by qPCR using primers for mitochondrial (*16S rRNA*) and nuclear (*CDKN2A/B*) gene. Males (left) and females (right) were fed either control food (black), or food supplemented with E2 (pink). Each symbol represents an individual fish. Bars represent minimum and maximum, and significance was measured using t-test.

**Figure S7:**
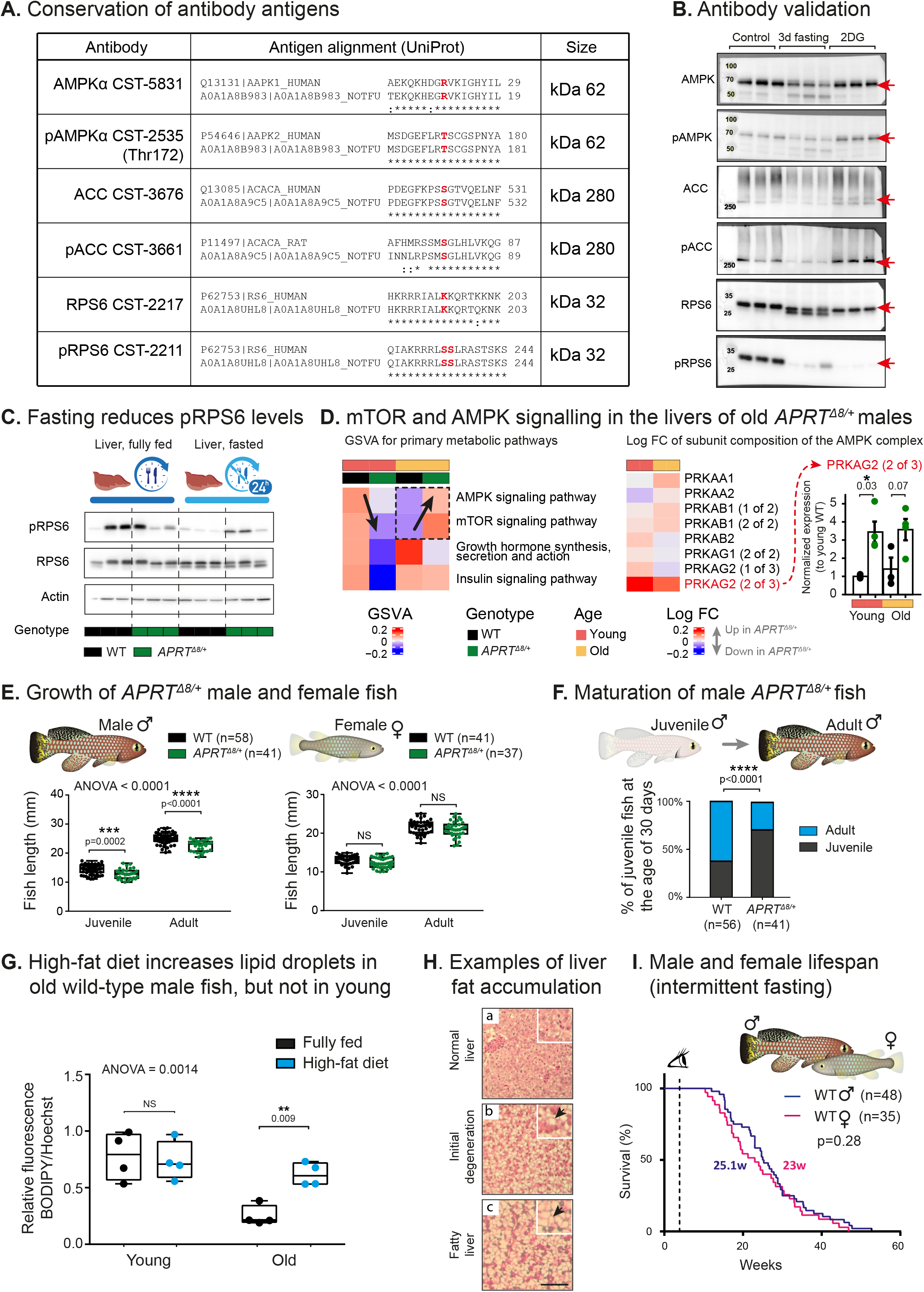
The effect of the *APRT* mutation on metabolic and physiological parameters. (**A**) Protein sequence conservation of the antigen sequence of the indicated antibodies and the corresponding killifish protein. (**B**) Validation of key antibodies used in this manuscript. Western blot of livers isolated from adult male fish under i) control condition; ii) chronic AMPK pathway activation via prolonged fasting (72 h fasting); iii) or acute AMPK pathway activation via immersion in 2-deoxy glucose (2DG, 20 mM for 3 h). Data are from 3 biological replicates for each experimental condition. (**C**) Western blot of livers isolated from old WT and *APRT^Δ8/+^* male fish. Experiments were performed under either control or 24 h fasting conditions. Data are from 3 biological replicates for each experimental condition. (**D**) Left: heatmap for transcriptional changes in fully-fed male livers using GSVA, highlighting primary metabolic pathways (KEGG). Arrows indicate overall increase or decrease in GSVA score. Right: Heatmap for gene expression changes (log2 fold change) between *APRT^Δ8/+^* and WT, for detected AMPK individual subunits and their respective isoforms. Additionally, upregulation of *PRKAG2* is also plotted as normalized expression values. Significance for differentially expressed genes was called by edgeR. (**E**) Measurement of growth rates during fish maturity. Experimental groups differ by sex (males or females), age (14 dph juveniles or 30 dph adults), and genotype (WT or *APRT^Δ8/+^*). Number of replicates for each experimental group are indicated (the same animals were measures as juveniles and adults). Bars represent minimum and maximum, and significance was measured by one-way ANOVA and Sidak’s correction for multiple tests. Exact p-values are indicated. (**F**) Scoring the onset of male maturity (using color changes as a proxy) in WT or *APRT^Δ8/+^* fish at different ages (juvenile or adults). Number of replicates for each experimental group are indicated. Significance was measured by χ^2^ test, and exact p-values are indicated. (**G**) Hepatic lipid accumulation and quantification in response to high-fat diet. Quantification of the relative fluorescence intensity of BODIPY/Hoechst (Fiji-ImageJ). Experimental groups differ by age (young or old) and feeding condition (control or high-fat diet). Bars represent minimum and maximum, and significance was measured by one-way ANOVA and Sidak’s correction for multiple tests. Exact p-values are indicated. (**H**) Histological characterization of livers in response to a short-term high-fat diet. **a**) Normal liver (controls). **b-c**) Pathology in response to high-fat diet (black arrows). First signs of degeneration are detected by fat accumulation in hepatocytes (b), or in more severe cases, by fatty liver (i.e. steatosis, c). Images were taken at the same magnification, scale bar 100 µm. Experiments were performed on old male fish (15 weeks, n = 3 for each feeding regimen). (**I**) Overlay of the male and female lifespans from WT fish under intermittent fasting conditions, using data from Figure 7d. p-values for differential survival in log-rank tests, median survival, and fish numbers are indicated. See Table S1 for additional information.

